# Early life exercise primes the neural epigenome to facilitate gene expression and hippocampal memory consolidation

**DOI:** 10.1101/2021.12.23.473936

**Authors:** AM Raus, TD Fuller, NE Nelson, DA Valientes, A Bayat, AS Ivy

**Author notes:** These authors contributed equally to the manuscript (AM Raus, TD Fuller, NE Nelson).

## Abstract

Aerobic exercise is well known to promote neuroplasticity and hippocampal memory. In the developing brain, early-life exercise (ELE) can lead to lasting improvements in hippocampal function, yet molecular mechanisms underlying this phenomenon have not been fully explored. In this study, adolescent transgenic mice harboring the “NuTRAP” (Nuclear tagging and Translating Ribosome Affinity Purification) cassette in Emx1 expressing neurons (“Emx1-NuTRAP” mice) undergo ELE followed by a hippocampal learning task, in order to determine the molecular underpinnings of exercise contributing to improved hippocampal memory performance. We simultaneously isolate and sequence translating mRNA and nuclear chromatin from a single hippocampus in a cell-type specific manner (excitatory neurons), demonstrate validity of our new technical approach, and couple multi-omics sequencing data to evaluate histone modifications H4K8ac and H3K27me3 and their influence on gene expression after ELE. We then evaluate new gene expression – histone modification relationships specifically during hippocampal memory consolidation that may play a critical role in facilitated memory after ELE. Our data reveal novel candidate gene-histone modification interactions and implicate gene regulatory pathways involved in ELE’s impact on hippocampal learning and memory.

## Introduction

Environmental experiences engage epigenetic mechanisms to modulate gene expression and cell function in post-mitotic neurons^1,2^. Histone modifications and DNA methylation are particularly important for neuronal adaptation to environmental signals by altering transcription and synaptic function^3, 4^. Behavioral outputs such as stress susceptibility, reward seeking, and long-term memory have been shown to result from changes to chromatin accessibility and gene expression in neurons^5–9^. In addition to this, the neuronal chromatin landscape undergoes waves of epigenetic modifications as a function of brain maturation itself^10–12^. Postnatal periods of heightened sensitivity to environmental stimuli can lead to lasting changes to cellular function and may result from temporally specific epigenetic mechanisms in the developing brain^13, 14^. Whether gene regulatory mechanisms in postmitotic neurons are uniquely influenced by early-life experiences to inform long-term function is a question that is just beginning to be explored. Identifying epigenetic processes involved in modulating cell function, particularly during brain development, is critical for understanding how early-life experiences impact long-term behavioral outcomes.

Aerobic exercise enhances performance on cognitive tasks involving the hippocampus in both adult humans and animal models^15, 16^. The type, timing, and duration of exercise exposure matters with regard to whether it has a persistent impact on hippocampal function^17–19^. Findings in both adolescent and adult rodents implicate a role for histone modifying enzymes in the mechanisms of exercise-induced benefits to hippocampal memory. Both voluntary exercise or treatment with a HDAC3 inhibitor enable hippocampal memory after a subthreshold learning stimulus, increase brain-derived neurotrophic factor (BDNF), and promote acetylation of H4K8 at the BDNF promoter^20–22^. This suggests that exercise engages epigenetic regulatory mechanisms to promote plasticity. Adult exercise also opens a temporal window for persistent improvements to memory performance when a reactivating exercise exposure is introduced^23^, suggesting a “molecular memory” of the initial exercise^23^. Although the majority of these studies have been performed in adults, more recent work also demonstrates the effects of exercise on hippocampal memory and associated alterations in neurotrophic factor expression, synaptic plasticity, and neurogenesis are similar in juvenile and adolescent periods^17, 24–27^. Previous work in our lab showed that early-life exercise (ELE) for either one week (juvenile period; postnatal days (P) 21-27) or three weeks (juvenile-adolescence; P21-41) facilitated hippocampal long-term memory formation in response to a learning stimulus typically insufficient for forming long-term memory. This finding was associated with increased long-term potentiation (LTP) as well as modulations to synaptic physiology in hippocampal CA1^17^. Notably, the hippocampal memory effects of juvenile ELE persisted two weeks after exercise cessation, which is potentially longer than the effect of exercise on adult hippocampal function^23^. Taking these findings together, it is possible that exercise (whether in early life or adulthood) may “prime” hippocampal function for response to future experiences (such as future exercise bouts or hippocampal learning events). Epigenetic mechanisms are strong candidates for the priming effects of exercise: the epigenome could harbor a “molecular memory” of the exercise experience by readying the chromatin landscape for efficient gene expression, thereby modulating neuronal function and behavioral output^22^. The specific mechanisms underlying sustained electrophysiological and behavioral effects observed in our studies of ELE have not been assessed from the perspective of a potential molecular memory of ELE within the epigenome.

In this study we describe a new approach to simultaneously isolate neuron-specific chromatin and translating mRNA from a single hippocampal homogenate, in order to address our hypothesis that ELE engages the epigenome to contribute to improved hippocampal memory and synaptic plasticity. We identify unique neuronal gene expression programs resulting from ELE and pair this data with ELE-induced changes in H4K8ac or H3K27me3. By crossing an Emx1-Cre transgenic line with mice harboring the NuTRAP (<u>Nu</u>clear tagging and <u>T</u>ranslating <u>R</u>ibosome <u>A</u>ffinity <u>P</u>urification^28^) cassette (termed “Emx1-NuTRAP” mice), we isolate whole nuclei and translating mRNA from a single hippocampus by combining the first steps of the INTACT (Isolation of <u>N</u>uclei <u>TA</u>gged in specific <u>C</u>ell <u>T</u>ypes^29^) and TRAP (<u>T</u>ranslating <u>R</u>ibosome <u>A</u>ffinity <u>P</u>urification^30^) published protocols. We have named our technical approach “SIT” (<u>S</u>imultaneous INTACT & <u>T</u>RAP). This technique yields DNA and RNA from the same homogenate that is of high-quality and sufficient concentrations for downstream sequencing procedures in this study, TRAP-seq and CUT&RUN-seq. Using neuron-specific, paired transcriptomic and epigenomic bulk sequencing, this study uncovers hippocampal gene expression programs and associated histone modifications that are driven by ELE to promote plasticity during hippocampal memory consolidation. Importantly, these data reveal an epigenetic signature resulting from ELE that may form the basis of persistently enabled hippocampal function.

## Results

### “Emx1-NuTRAP” mouse allows for simultaneous isolation of nuclear chromatin and translating mRNA from a single population of hippocampal neurons

Our initial goal was to obtain translating mRNA and nuclear DNA from a single population of neurons to couple transcriptional network changes with alterations in histone modifications resulting from ELE. To do this, we developed a transgenic mouse line (Emx1-NuTRAP) and experimental protocol (SIT protocol). We crossed male NuTRAP reporter mice^28^ with female Emx1-Cre^31, 32^ mouse lines to generate Emx1-Cre; NuTRAP transgenic progeny for this study (“Emx1-NuTRAP”; Fig. 1A). Emx1-expressing neurons are predominately excitatory^31^ and are necessary for hippocampal neurogenesis and motor skill learning^33^. In the presence of a *loxP* site-flanked sequence, Emx1-Cre facilitates recombination in approximately 88% of neurons in the neocortex and hippocampus, and in less than 2% of GABAergic inhibitory interneurons in these regions^31^. Emx1-Cre also targets neural progenitor cells and can be expressed in mature astrocytes in striatum^34^. Of note, we exclusively used Emx1-Cre female mice in our breeding schemes given that the Cre-recombinase has been reported to be expressed in a subset of male germline cells^35^.

**Figure 1.**
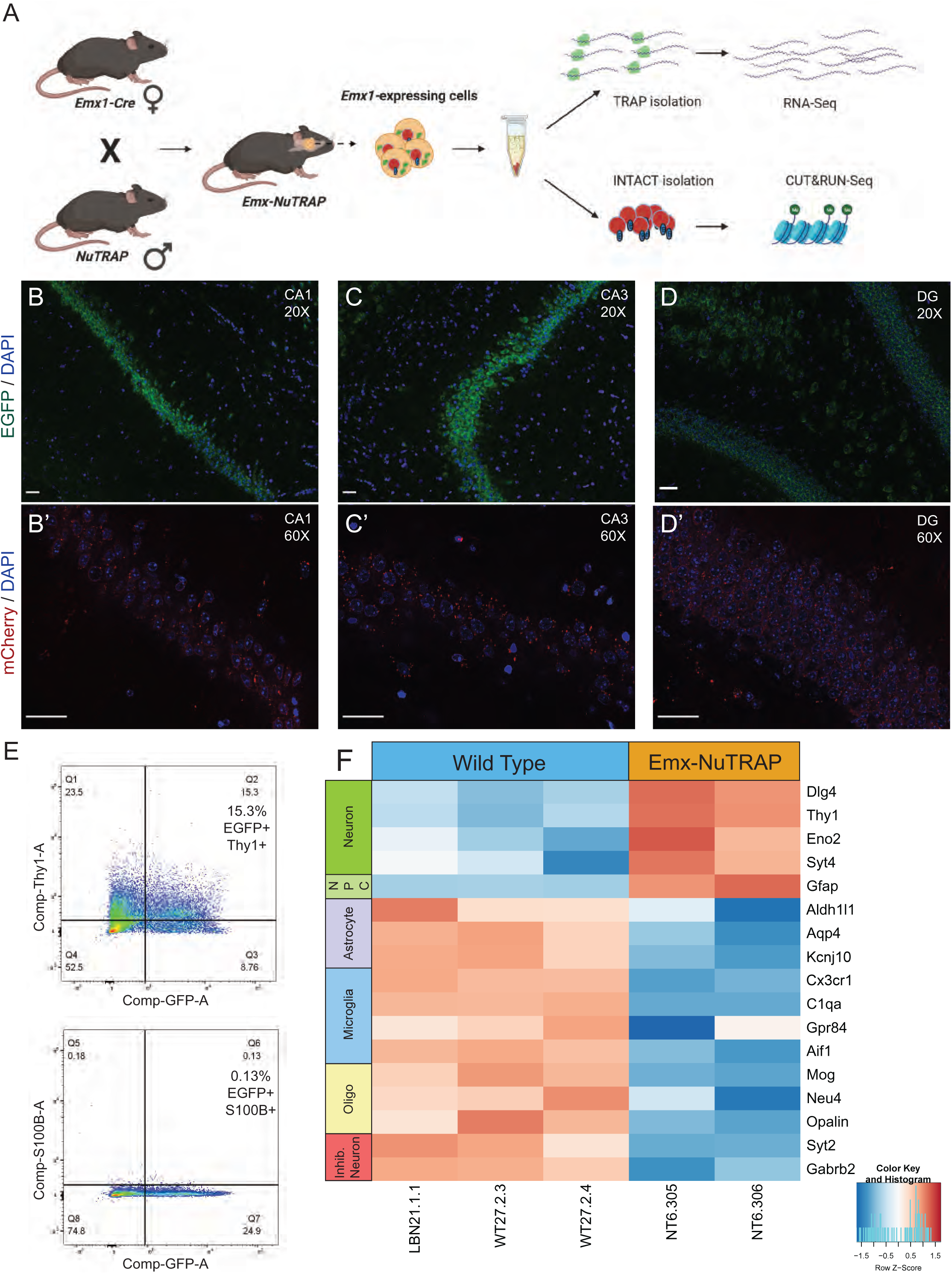
Generation and validation of the Emx1-NuTRAP mouse. (A) Schematic representation of the Emx1-NuTRAP mouse generation and workflow of the Simultaneous INTACT and TRAP (“SIT”) protocol. (B) Immunofluorescence imaging of the CA1 region of the hippocampus at 20x objective and (B’) 60x objective after incubation with an mCherry antibody overnight. (C) Immunofluorescence imaging of the CA3 and CA2 region of the hippocampus at 20x objective and (C’) 60x objective after incubation with an mCherry antibody overnight. (D) Immunofluorescence imaging of the DG region of the hippocampus at 20x objective and (D’) 60x objective after incubation with an mCherry antibody overnight. Scale bars are set to 50µm for all images (B-D). (E) Flow cytometry for EGFP, THY-1, and S100β (15.3% of cells THY-1+/EGFP+ and 0.13% of cells S100β+/EGFP+). (F) Heatmap of differentially expressed neuronal and other cell type markers from RNA-seq data comparing TRAP-isolated RNA from our Emx1-NuTRAP mice and previously generated dorsal hippocampal tissue RNA from wild type mice.

We sought to validate neuron-specific expression of the NuTRAP cassette. First, immunohistochemistry of Emx1-NuTRAP hippocampal slices show distinct GFP and mCherry expression in CA1 and CA3 pyramidal neurons (Fig. 1B-C) as well as the granule cell layer of the dentate gyrus (DG; Fig. 1D). Furthermore, mCherry positive nuclei remained intact and bound to the magnetic beads after our modified nuclear isolation procedure (Supplementary Fig. 1A). As Emx1-expressing neural stem cells can become astrocytes^31^, we sought to determine whether a significant number of mature astrocytes were obtained in the population of isolated cells. Flow cytometry experiments using THY-1 as a neuronal marker and S100β as an astrocytic marker revealed a distinct population of cells double positive for GFP and THY-1, whereas a S100β and GFP double positive cell population was absent (Fig. 1E and Supplementary Table 1).

To perform <u>S</u>imultaneous INTACT & <u>T</u>RAP (“SIT”), hippocampal tissue was dissected from both brain hemispheres, combined, and homogenized in one sample tube (Fig. 1A). We then performed the “SIT” method on isolated samples by starting the protocol with the beginning steps from the INTACT procedure modified to include cycloheximide. Cycloheximide works rapidly to inhibit protein synthesis and is used for maintaining crosslinks between translating mRNA and ribosomal subunits during purification in the traditional TRAP method. Despite the presence of cycloheximide, nuclear morphology from Emx1-expressing neurons is overall unchanged (Supplementary Fig. 1A). Additionally, a Bioanalyzer was used to determine if there were cycloheximide-induced double-stranded DNA breaks in our nuclear preparation that could interfere with downstream DNA sequencing applications (such as ATAC-, CUT&RUN-, or CUT&Tag-seq). We found no evidence of DNA double strand breaks generating fragments of less than 1kb (Supplementary Fig. 1B). Following magnetic purification of biotin-labeled nuclei, the supernatant fraction was removed and taken through TRAP, while the pelleted nuclei were processed through the remaining steps of INTACT (see Methods). Using qPCR, we found that TRAP isolated mRNA had significantly reduced expression of *Mog* and *Cd11b* compared to total RNA, suggesting depletion of oligodendrocyte and microglial populations in TRAP mRNA (Supplementary Fig. 1C). Combining bilateral hippocampi from a single mouse yielded TRAP-isolated mRNA of high quality and sufficient concentration for sequencing (RIN > 8 for all samples, average yield RNA = 14.367 ng/ul; Supplementary Table 2 and Supplementary Fig. 1D). INTACT-isolated nuclei were further processed using the CUT&RUN (<u>C</u>leavage <u>U</u>nder <u>T</u>argets and <u>R</u>elease <u>U</u>sing <u>N</u>uclease^36^) method to isolate antibody-specific protein-DNA interactions for downstream DNA sequencing. The resulting DNA libraries were of high quality and concentration when using specific antibodies (H4K8ac: average size = 1238 bp, average concentration = 126.8 nM; H3K27me3: average size = 1032 bp, average concentration = 146.5 nM; Supplementary Table 2). In contrast, the resulting DNA libraries using the non-specific IgG control had substantially lower concentrations (IgG: average size = 1035 bp, average concentration = 24.2 nM; Supplementary Table 2) further indicating that nuclear DNA from both isolations was of high starting quality.

To confirm our TRAP-isolated hippocampal mRNA came primarily from excitatory neurons, we compared RNA-seq data from whole dorsal hippocampal tissue of wild type mice to our TRAP-isolated RNA-seq data (TRAP-seq) from Emx1-NuTRAP mice to assess for neuronal gene enrichment. We found that TRAP-isolated mRNA had significant enrichment of several neuronal genes, including *Dlg4*, *Thy1*, *Eno2*, and *Syt4*, as well as a significant reduction in astrocytic, microglial, oligodendrocyte, and inhibitory neuronal genes (Fig. 1F). There was a relative expression increase of glial fibrillary astrocytic protein (GFAP) in our TRAP samples. This gene can also be expressed in neural stem populations that were present in our whole hippocampus samples, so this finding may be a reflection of the neural stem cell population of the dentate gyrus^37, 38^. Overall, these findings suggest that the Emx1-NuTRAP mouse model is a valid tool for neuron-enriched isolation of sequencing-grade, translating mRNA and nuclear chromatin from a single brain tissue homogenate.

### Performing INTACT/TRAP on the same or separate cell suspensions yields highly comparable TRAP-seq and CUT&RUN-seq results

Previous methods for performing INTACT and TRAP using NuTRAP mice have taken separate tissue homogenates for each procedure^28, 39^. In this study, we performed SIT on a single tissue homogenate containing bilateral hippocampi (the products of SIT are referred to as “simultaneous isolations”). To determine if our approach of SIT is comparable to separate INTACT and TRAP procedures, we also performed separate INTACT and TRAP from hippocampal tissue obtained from one brain hemisphere for each method, counterbalancing for left versus right hemispheres (we refer to this protocol as “separate isolations”). A Bioanalyzer was used to determine the amount and quality of the RNA obtained from each type of isolation (Separate isolations: average RNA concentration = 7.805 ng/ul, average RIN = 9.5; Simultaneous isolations: average RNA concentration = 14.367 ng/ul, average RIN = 9.3; Supplementary Table 2). The average RNA yield from the separate isolations (using a unilateral hippocampus) was approximately equal to half of the average yield of the simultaneous isolations (which combined bilateral hippocampi; Supplementary Table 2). Similarly, the final library concentrations for the separately isolated IgG CUT&RUN-seq libraries were also approximately half the concentration of the simultaneous isolations (average simultaneous: 24.2nM, average separate: 11nM; Supplementary Table 2). We interpret this to mean that nuclear DNA was fully intact in the simultaneous isolation because we did not obtain substantially more than double the concentration in the simultaneous vs separate isolations. Unilateral hippocampal homogenates yielded sufficient sequencing concentrations and quality to allow for library preparations from individual mice (Supplementary Table 2).

To determine if normalized TRAP-seq counts were similar between the two isolation methods, we performed a Spearman’s correlation between the datasets from sedentary mice. The two TRAP-seq datasets were found to be highly correlated, with R values greater than 0.5 and p value less than 2.2 x 10^-16^ (R=1, p<2.2×10^-16^; Fig. 2A). CUT&RUN-seq was used to identify genomic regions interacting with either H4K8ac, an activating histone post-translational modification (PTM), or H3K27me3, a generally repressive histone PTM. We again applied Spearman’s correlation to understand whether the simultaneous vs separate INTACT isolation methods could influence CUT&RUN-seq peak distribution. We compared normalized count data for CUT&RUN-seq peaks across a representative chromosome (chromosome 2). We binned 100bp increments along the entire chromosome from the simultaneous and separate isolations using datasets generated from sedentary mice. Normalized sequencing counts, which reflected reads assigned to binned genomic regions along chromosome 2, were highly similar between conditions (H4K8ac: R=0.65, p<2.2×10^-16^; H3K27me3: R=0.81, p<2.2×10^-16^; Fig. 2B-C). Taken together, these experiments suggest that translating mRNA and nuclear DNA isolated from hippocampal homogenates using either simultaneous or separate isolation procedures are comparable in terms of quality, concentration, functional characterization, and normalized sequencing reads.

**Figure 2.**
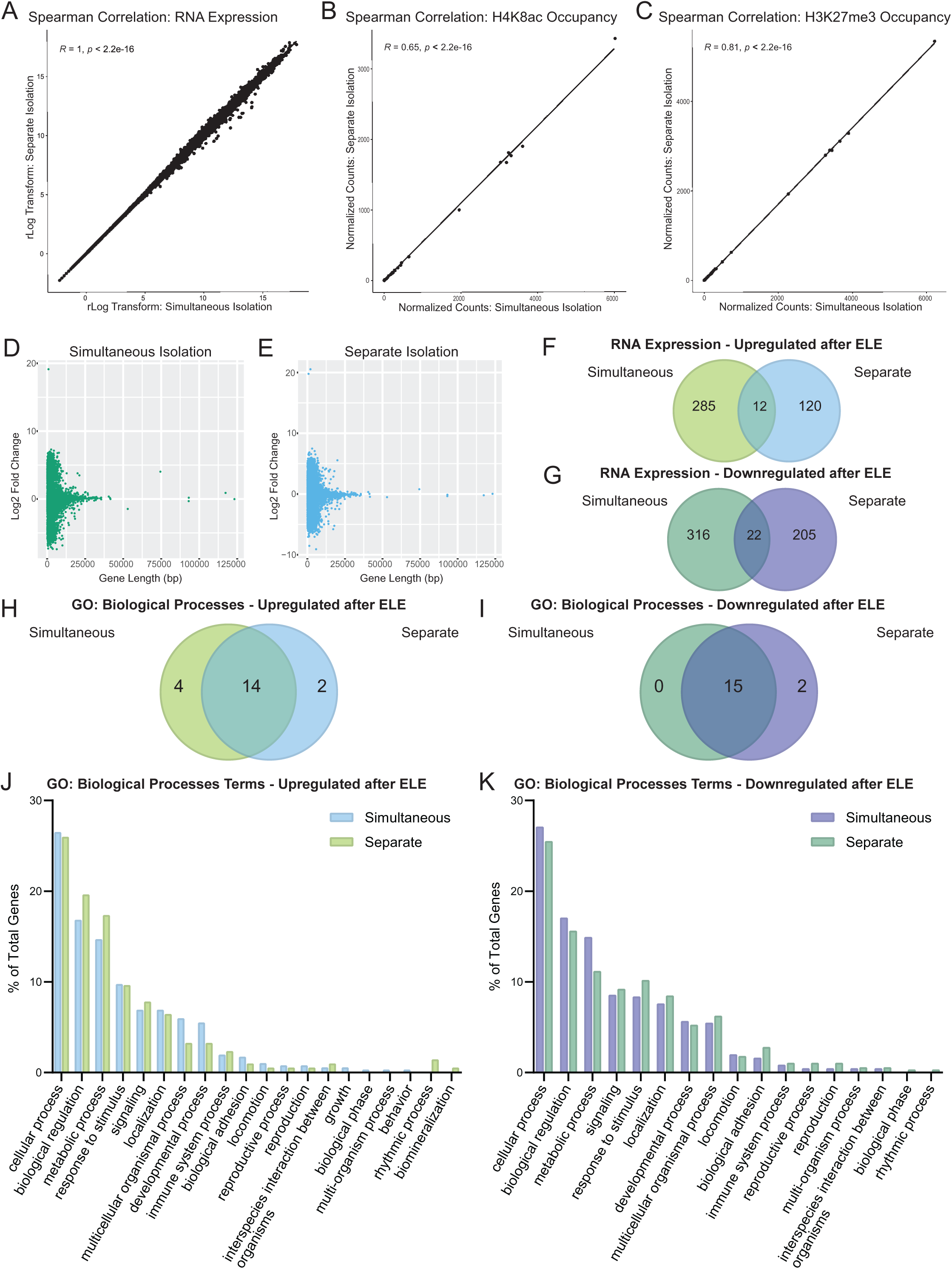
Simultaneous and separate isolations of DNA and RNA from Emx1-NuTRAP mice are comparable. For Fig. 2A-E, n=2 mice for simultaneous isolations and n=3 mice for separate isolations. (A) Spearman’s correlation for RNA expression between separately isolated RNA and simultaneously isolated RNA (R=1, p<2.2×10^-16^). (B) Spearman’s correlation between separate and simultaneously isolated DNA from CUT&RUN-seq for H4K8ac with 100bp chromosome 2 position bins (R=0.65, p<2.2×10^-16^). (C) Spearman’s correlation between separate and simultaneously isolated DNA from CUT&RUN-seq for H3K27me3 with 100bp chromosome 2 position bins (R=0.81, p<2.2×10^-16^). (D) Gene length versus fold change distribution plot for simultaneously isolated RNA. (E) Gene length versus fold change distribution plot for separately isolated RNA. (F) Genes upregulated in TRAP-seq after ELE in either simultaneous (n=2 sedentary mice and n=3 ELE mice) or separate isolations (n=3 mice per group). (G) Genes downregulated in TRAP-seq after ELE in either simultaneous or separate isolations. (H, I) Panther Gene Ontology: Biological Processes Venn diagram for genes upregulated (H) or downregulated (I) after ELE in either separate or simultaneous isolations. (J, K) Panther Gene Ontology: Biological Processes bar graph of gene distributions for genes upregulated (J) and downregulated (K) after ELE in either separate or simultaneous isolations.

We next wanted to determine if the different isolation methods could bias resulting gene expression on the basis of gene length. When plotting gene length against log fold change of gene expression, we see a similar distribution pattern of ELE-induced differentially expressed hippocampal genes (DEGs) between the simultaneous and separate methods (Fig. 2D-E). We then compared the biological categories of the ELE-induced DEGs obtained using either the separate or the simultaneous isolation methods using Panther Gene Ontology (GO). Although many of the genes did not overlap (Fig. 2F-G and Supplementary Table 3), most GO terms did (14/20 GO terms for upregulated genes and 15/17 for downregulated genes; Fig. 2H-I and Supplementary Table 3). Furthermore, there was high similarity between the percentage of genes found in each of the GO categories (Fig. 2J-K). Taken together, our results demonstrate that the simultaneous approach for isolating and sequencing mRNA and nuclear chromatin can be performed using a single hippocampal homogenate, and can generate highly similar results as more traditional methods using separate cell samples to pair different types of sequencing results.

### Neuron-specific gene expression and histone PTMs are functionally comparable between left and right hippocampal hemispheres

Many studies take advantage of the brain’s structural symmetry by using tissue from each hemisphere for separate molecular processing. Prior evidence demonstrates that hippocampal lateralization can influence LTP, hemisphere-specific glutamate receptor density, and performance in certain memory tasks^40^. We wanted to determine if there were differences in transcriptional programs and biological processes resulting from ELE in left vs right hippocampi. Using the separate isolation approach described above, we compared ELE-induced differential gene expression between left and right hemispheres. We found that although individual gene expression patterns were different between hippocampi originating in the left and right brain hemispheres (Fig. 3A-B and Supplementary Table 4), the Panther GO: Biological Processes terms were similar, with most categories overlapping (Fig. 3C-F and Supplementary Table 4). This could suggest spatial or hemispheric assignment of genes with functional similarity.

**Figure 3:**
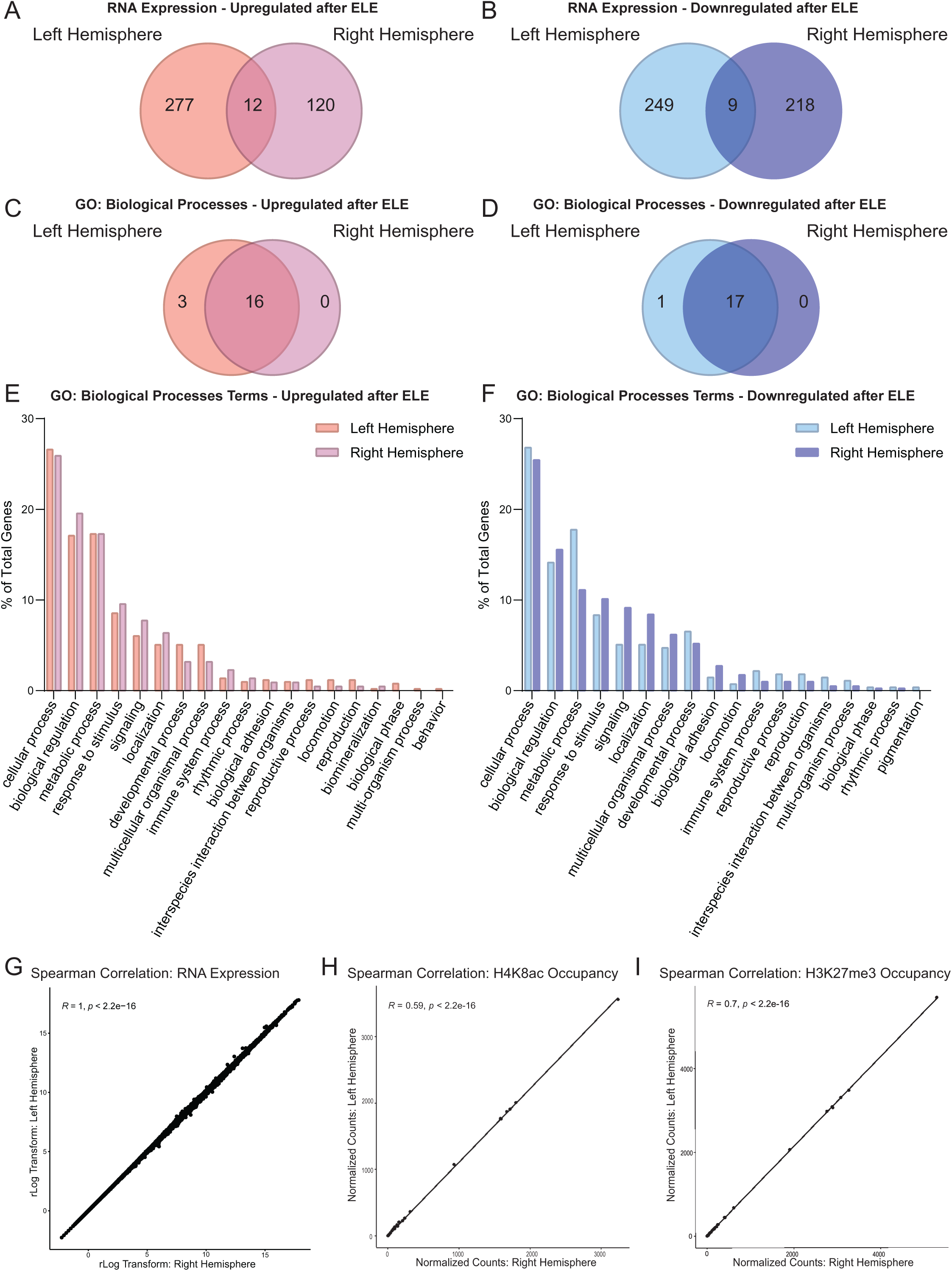
Histone modifications and functional categorizations of gene expression are similar regardless of hemispheric origin of hippocampal tissue. (A-B) Venn diagram of genes upregulated (A) or downregulated (B) by ELE identified between the left and right hemispheres by the separate isolation protocol TRAP-seq (n=3 mice per group). (C-D) Venn diagram of Panther Gene Ontology: Biological Process terms for genes upregulated (C) or downregulated (D) by ELE identified between the left and right hemispheres by the separate isolation protocol TRAP-seq. (E-F) Percent of genes fitting into each GO category for the left and right hemisphere for genes upregulated (E) and downregulated (F) by ELE by separate isolation TRAP-seq. (G) Spearman’s correlation between the left and right hemispheres TRAP-seq data from the separate isolation protocol of sedentary rlog normalized expression (R=1, p<2.2×10^-16^). (H) Spearman’s correlation between hemispheres for CUT&RUN-seq for H4K8ac normalized count data for chromosome 2 in 100bp bins (R=0.59, p<2.2×10^-16^). (I) Spearman’s correlation between hemispheres for CUT&RUN-seq for H3K27me3 normalized count data for chromosome 2 in 100bp bins (R=0.7, p<2.2×10^-16^).

Finally, we wanted to determine if left and right hippocampal hemispheres had significant differences in DEGs and CUT&RUN-seq peak distributions at baseline (without exercise). We performed Spearman’s correlation on the transcript counts from the TRAP-seq data (R=1, p<2.2×10^-16^; Fig. 3G), and the CUT&RUN-seq 100 bp binned peak counts along chromosome 2 for H4K8ac (R=0.59, p<2.2×10^-16^; Fig. 3H) and for H3K27me3 (R=0.70, p<2.2×10^-16^; Fig. 3I). We found that the seq peaks and transcript counts were highly similar with significance considered as R values greater than 0.5 and p value less than 2.2 x 10^-16^ (Fig. 3G-I). Overall, left and right hippocampal hemispheres did not demonstrate significant differences in normalized sequencing counts and peak distributions in the sedentary condition, suggesting that choice of hippocampal hemisphere is a less important factor to consider in obtaining representative data on transcription and transcriptional regulation via histone modifications.

### Early-life exercise promotes expression of plasticity-related genes and is associated with transcriptional regulatory pathways implicated in hippocampal memory

To identify the transcriptional effects of ELE in hippocampal neurons, we performed cell-type specific bulk sequencing on TRAP-isolated mRNA extracted from Emx1-NuTRAP hippocampi using our “SIT” protocol. We compared DEGs from sedentary mice vs ELE (running distances shown in Supplementary Fig. 2). Using DESeq2 to assess for DEGs (>30% expression increase with a p value < 0.05), we found that ELE alters gene expression in Emx1 expressing neurons (297 upregulated and 338 downregulated genes; Fig. 4A and Supplementary Table 3). Many of the genes upregulated after ELE are known to be involved in exercise and/or hippocampal memory mechanisms, including *Bdnf* ^41^ and *Nr4a1*^42^. To functionally categorize ELE-induced DEGs, we performed a Panther Gene Ontology (GO) analysis^43^ focusing on the Molecular Function categorization and separated by upregulated and downregulated genes. Regardless of gene expression directionality, GO term categories with the most genes functionally assigned to them were “binding”, “catalytic activity”, “molecular function regulator”, “transporter activity”, “molecular transducer activity”, and “structural molecule activity” (Fig. 4B). Many of the upregulated genes driving these categories are known to have critical roles in neuronal function (*Kcna1, Slc24a4, Stxbp5l, Gabra2,* and *Camk2n2*), neurodevelopment (*Artn, Kdm7a, Sox21, Gap43,* and *Efna5*), and hippocampal memory (*Bdnf, Nr4a1* and *Dusp5*).

**Figure 4:**
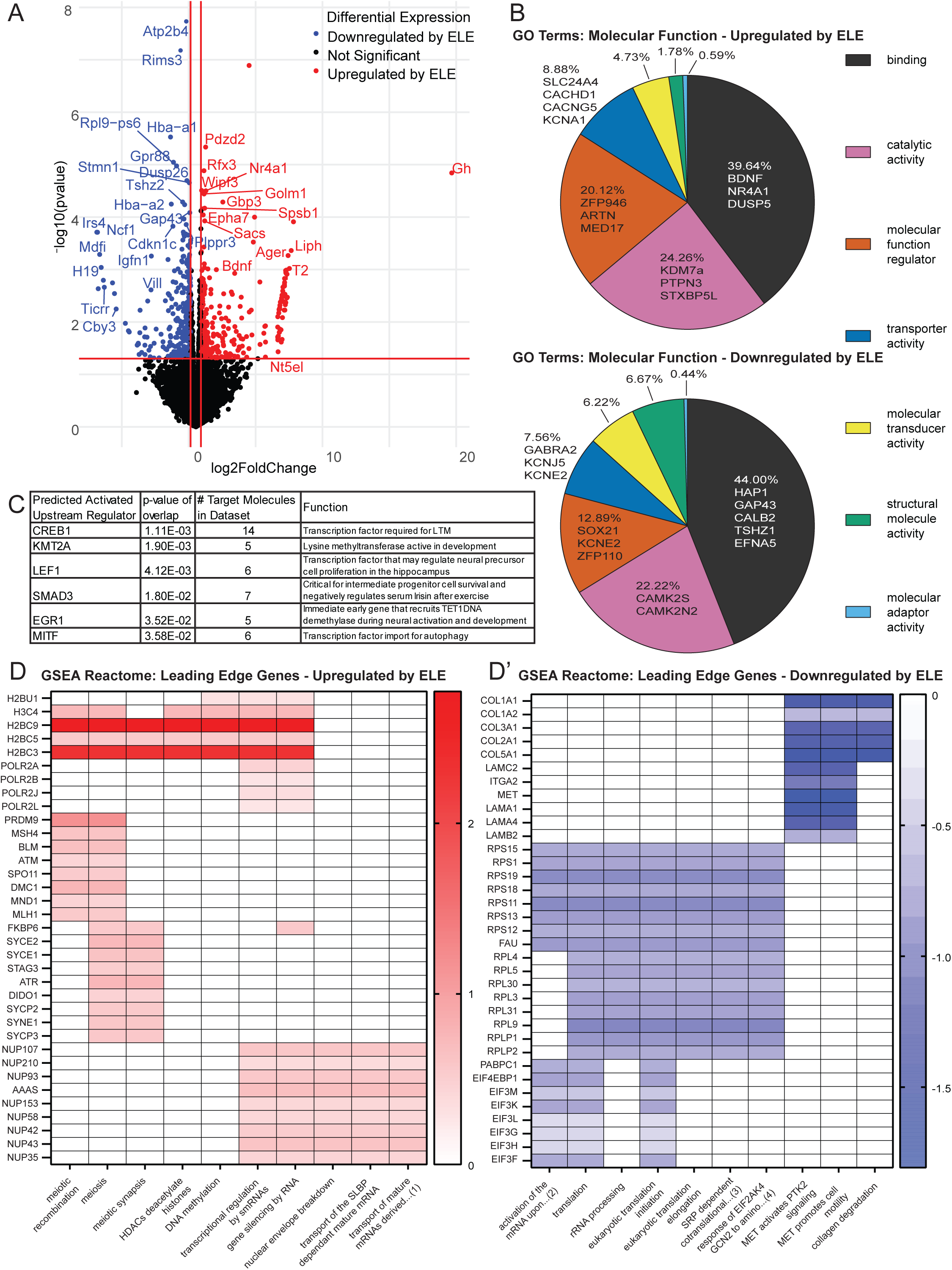
ELE leads to transcriptomic changes in hippocampal neurons during adolescence. (A) Volcano plot of differentially expressed genes reaching significance identified by TRAP-seq on ELE versus sedentary (n=2 sedentary mice and n=3 ELE mice, absolute value log_2_ fold change > 0.3785 and p-value < 0.05). (B) Panther Gene Ontology: Molecular Function top terms by most genes assigned. (C) Top 6 “Upstream Regulators” identified by Ingenuity Pathway Analysis (IPA). (D) Representative Gene Set Enrichment Analysis: Reactome leading-edge diagrams showing genes upregulated (D) or downregulated (D’) in ELE, and their categories of enrichment. *Abbreviated terms in (D): (1) “transport of mature mRNAs derived from intronless transcripts”, (D’): (2) “activation of the mRNA upon binding of the cap binding complex and EIFs and subsequent binding to 43S”, (3) srp dependent cotranslational protein targeting to membrane, and (4) response of eif2ak4 gcn2 to amino acid deficiency.

To evaluate possible transcription factors and upstream regulators implicated by ELE-induced activated gene networks, we applied Qiagen’s Ingenuity Pathway Analysis (IPA) to our TRAP-seq dataset^44^. We identified significant canonical signaling pathways implicated in ELE effects on hippocampal neuronal function (Supplementary Fig. 3 and Supplementary Table 5). Additionally, the top six upstream regulators by significance included CREB1, KMT2A, LEF1, SMAD3, EGR1, and MITF. Each of the top five transcription factors have been implicated in hippocampal development or function (Fig. 4C and Supplementary Table 5)^45–52^. The transcription factors KMT2A and LEF1 have not been previously associated with the effects of exercise on hippocampal function. Interestingly, CREB1 (through its associated CBP^53^) and KMT2A (through its methyltransferase activity^46^) both have histone modifying properties. LEF1 has been shown to regulate neural precursor proliferation in the hippocampus^47^. SMAD3 is critical for intermediate progenitor cell survival and negatively regulates serum insulin after exercise^48, 49^. EGR1 is an immediate early gene that recruits TET1 DNA demethylase during neural activation and development^50^. MITF was the sixth upstream regulator identified and is a transcription factor linked to autophagy mechanisms^51^. These specific transcription factors were not significantly differentially expressed in our TRAP-seq dataset; however, their activity may not be linked to a change in their own expression after ELE but rather modulated by exercise to influence programs of gene expression.

To further understand whether the transcriptional profiles resulting from ELE were enriched for specific *a priori* assigned molecular functions, we performed Gene Set Enrichment Analysis (GSEA)^54^. We evaluated the “Reactome” category of functional gene sets followed by a leading-edge analysis to further determine which genes were driving the significant categories (Fig. 4D and Supplementary Table 6). Of the DEGs upregulated after ELE, several interesting categorizations and the genes driving them were revealed. The genes *H3c4*, *H2bc3*, *H2bc5*, and *H2bc9* genes, which encode histone family member proteins, were driving the categories: “DNA methylation”, “HDACs deacetylate histones”, and “transcriptional regulation by smRNAs”. “Meiosis” and “meiotic recombination” also emerged in these results which was unusual; however, leading edge analysis showed many of these genes (*Atm, Blm, Msh4*, and *Mnd1*) to be generally involved in cell-cycling processes. Exercise is well known to increase adult neurogenesis in hippocampal dentate gyrus and can explain cell cycle gene enrichment in our dataset^16^. Several nucleoporin complex genes (*Nup35/ 42/ 43/ 50/ 58/ 62/ 93/ 107/ 153/ 210*) were also identified for their associations with categories such as: “gene silencing by RNA”, “nuclear envelope breakdown”, and “transport of mature mRNAs derived from intronless transcripts”. *Nup* genes form a variety of nuclear pore complexes that play critical roles in cellular processes including cell-cycle regulation, cellular differentiation, and epigenetic control^55^.

Of the downregulated pathways identified, translation-associated categories (“translation”, “eukaryotic translation initiation/elongation”, and “SRP-dependent co-translational protein targeting to membrane”) were driven by significant downregulation of ribosomal protein gene families (*Rps*, *Rpl*, *Rplp*) and eukaryotic initiation factor 3 (*eIF3*) subunits. Over-expression of *eIF3* subunits has been linked to neurodegeneration, and altered expression of *eIF3* has been associated with neurodevelopmental disorders^56^. Additionally, collagen genes (*Col1a1*, *Col1a2*, *Col2a1*, *Col3a1*, and *Col5a1*) were downregulated after ELE, leading to the identification of categories including: “MET activates PTK2 signaling”, “MET promotes cell motility”, and “collagen degradation”. *Col1a1* and *Col1a2* have been previously identified as putative aging genes that decrease in their expression after chronic exercise in female adult rodents^57^. By analyzing these enriched pathways and the networks of genes driving them, we were able to identify several expected, as well as unexpected, transcriptional programs activated by ELE that could be unique to exercise timing during the juvenile-adolescent period.

### Neuronal H4K8ac is enriched and H3K27me3 is reduced at a subset of plasticity genes after ELE

The power of the Emx1-NuTRAP mouse model (coupled with the “SIT” technical approach) is the ability to directly pair differential gene expression with governing epigenetic mechanisms. This is achieved by obtaining both TRAP-seq and epigenomic-sequencing data from the same cell population (Fig. 1A). To investigate ELE-induced changes in histone PTMs across the genome, hippocampal nuclei were obtained from ELE and sedentary mice using the INTACT method followed by CUT&RUN-seq for the modifications H3K27me3 and H4K8ac from nuclear chromatin. We chose these two histone PTMs for their previously described functions. H4K8ac is a permissive modification that is enriched at *Bdnf* after adult exercise, and its presence correlates with improved hippocampal memory^22^. H3K27me3 is a repressive histone mark and a common control in CUT&RUN-seq studies^36, 58^ and is decreased at the *Bdnf* promoter region after contextual fear conditioning training^59^. Peaks were called using SEACR^60^ and histone PTM enrichment peaks were overlapped with ELE-induced DEGs obtained from our TRAP-seq experiments described above (Fig. 5A and A’). We evaluated for the presence of H4K8ac and H3K27me3 peaks in union with upregulated and downregulated mRNA as a result of ELE. 93 upregulated genes had new H4K8ac peaks, while 35 downregulated genes had new H3K27me3 peaks (Fig. 5B and Supplementary Table 7). Notably, of those 93 upregulated genes with H4K8ac peaks, 14 also had new H3K27me3 peaks as a result of ELE. 17 genes had presence of H3K27me3 in both conditions (ELE and sedentary; Fig. 5A). We interpret these results to mean that transcription of these 93 genes was promoted as a result of ELE-induced H4K8ac.

**Figure 5:**
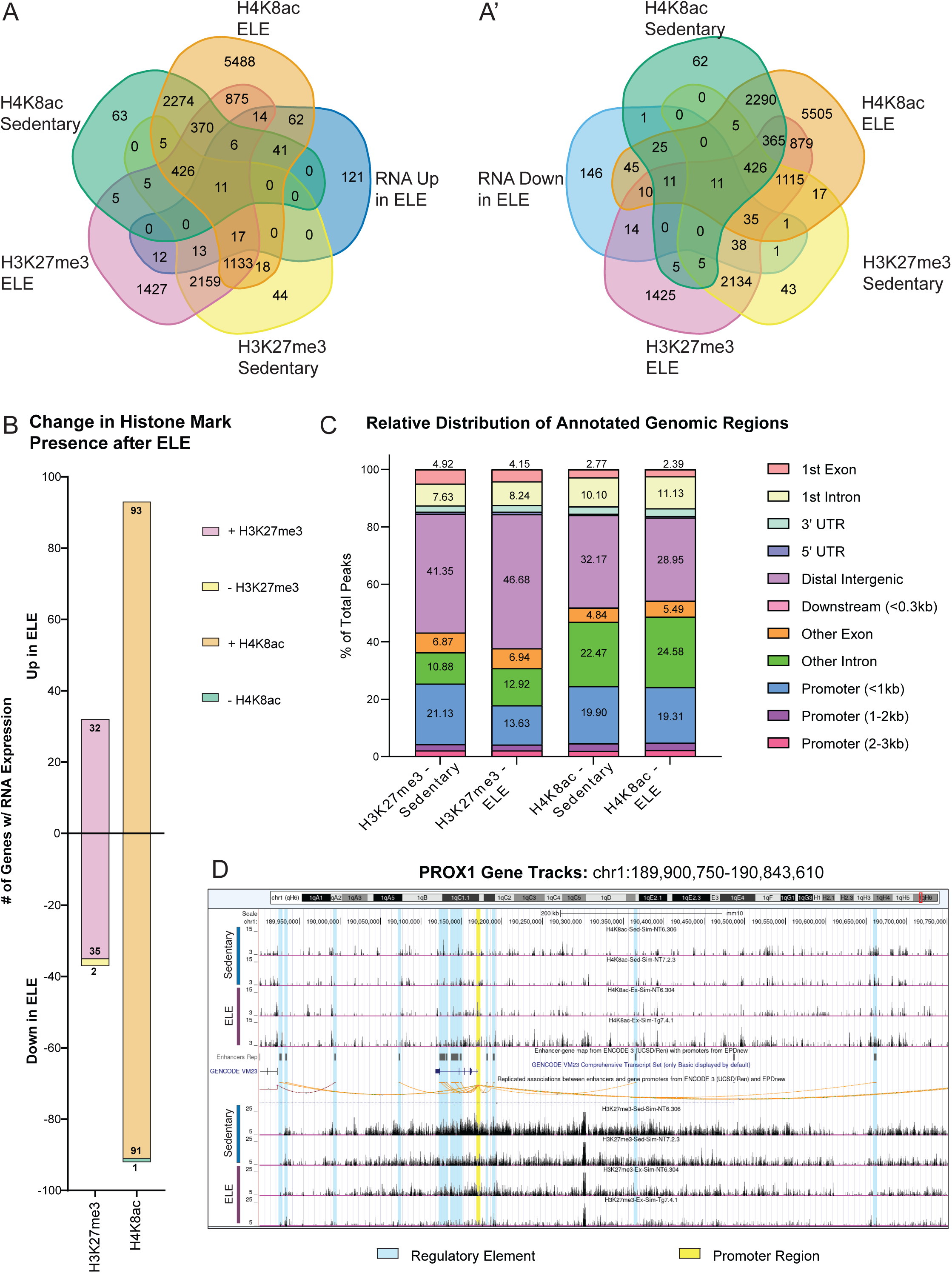
ELE alters H4K8ac and H3K27me3 occupancy. (A) Venn diagram of gene peaks identified in CUT&RUN-seq for H4K8ac and H3K27me3 by SEACR (top 1% of peaks FDR<0.1) overlapped with genes upregulated (A) and downregulated (A’) in RNA by ELE (n=2 sedentary mice and n=3 ELE mice). (B) CUT&RUN-seq SEACR peak calls at genes that overlap with genes differentially expressed by ELE as determined by TRAP-seq. (C) Stacked bar of the relative distribution of peak calls at various genomic regions identified by CUT&RUN-seq called using SEACR. (D) Representative track of CUT&RUN-seq data around *Prox1* showing differential peak height between ELE and sedentary conditions for H4K8ac (top) and H3K27me3 (bottom).

The loss of either H3K27me3 or H4K8ac after ELE did not substantially correlate with changes in translating mRNA expression (Fig. 5B). This result may indicate that ELE-induced differential gene expression is associated with the addition, rather than removal, of these two histone modifications. We found one gene that was downregulated and had loss of H4K8ac (*Asphd1*), and two genes downregulated with loss of H3K27me3 (*Fdxr* and *Rtl1*; Fig. 5B and Supplementary Table 7). Of particular interest, reduced expression of paternally-inherited *Rtl1* is associated with improved hippocampal memory performance, whereas overexpression results in memory deficits^61, 62^. Unsurprisingly, the majority of histone PTM peaks newly present as a result of ELE did not correlate with gene expression changes. However, notable upregulated genes with new H4K8ac after ELE include *Prox1*, *Sntb2*, and *Kptn*. PROX1 is involved governing differentiation, maturation, and DG versus CA3 cell identity in intermediate progenitor cells of the hippocampal DG^63^, while SNTB2 is involved in G protein-coupled receptor cell signaling^64^. KPTN encodes a protein involved in cytoskeletal cell structure, and its mutation can cause neurodevelopmental disability and seizures^65^. Interestingly, downregulated genes found to have new H3K27me3 after ELE included *Col3a1*, *Efnb2*, *Epop*, and *Myoc*. COL3A1 and MYOC are structural proteins involved in the extracellular matrix and the cytoskeleton respectively^66–68^. EFNB2 is a signaling molecule involved in cell migration and its haploinsufficiency can cause neurodevelopmental disability^69–71^. EPOP is a known editor of the chromatin landscape by altering H2Bub and H3K4me3 distributions^72, 73^.

Although changes in gene expression associated with H3K27me3 peaks were less numerous than those associated with H4K8ac, we noticed a striking difference in the distribution of peaks across the genome (Fig. 5C). After ELE, the distribution of H3K27me3 decreases in the promoter region and increases in the distal intergenic and intronic regions (Fig. 5C). To investigate how ELE-induced changes to histone PTM distribution might alter gene expression, we looked at the distribution of peaks around *Prox1* (Fig. 5D). Most apparent changes in histone PTM peaks occurred in regions annotated as regulatory regions, rather than the promoter region, which is concordant with our data showing that most histone PTM changes happen outside of the promoter region. Taken together, these results suggest that ELE promotes the new addition of H4K8ac and H3K27me3 PTMs to regulate gene expression. Given that most identified peaks did not correlate with DEGs, they may instead implicate regions of interest that are “primed” for facilitated gene expression following future stimuli, such as learning.

### ELE alters H4K8ac and H3K27me3 occupancy at regulatory regions of genes involved in hippocampal memory consolidation

Prior work from our lab and others has found that both ELE and adult exercise can facilitate hippocampal long-term memory formation in mice exposed to a typically sub-threshold learning event (3 minutes of object location memory (OLM) training, which is normally insufficient for LTM formation in sedentary mice)^17, 22^. These findings suggest that ELE may “prime” neuronal function to facilitate hippocampal learning. In this experiment, we investigate whether ELE-enabled hippocampal memory is associated with altered presence of histone PTMs H4K8ac and H3K27me3 in regulatory regions of genes associated with learning and hippocampal plasticity; the prediction being that alterations of histone PTM occupancy as a result of ELE could promote gene expression programs necessary for facilitating memory consolidation. Wild type animals underwent ELE followed by OLM training on P42 and were sacrificed 60 minutes later (Fig. 6A). Dorsal hippocampi were dissected, and tissue was homogenized and processed for bulk RNA-seq. These data were compared to the ELE-induced DEGs, H4K8ac, and H3K27me3 occupancy occurring without learning (generated from Emx1-NuTRAP mice TRAP-seq and CUT&RUN-seq). We categorized DEGs based on the experimental group of wild type animals: ELE alone, ELE+3 min learning stimulus, Sedentary+3 min learning stimulus, and Sedentary+10 min learning stimulus. To discover ELE-induced genes “primed” for regulation during memory consolidation, we focused specifically on <u>c</u>andidate <u>g</u>enes <u>a</u>ssociated with <u>m</u>emory after <u>e</u>xercise (cGAME), <u>c</u>andidate <u>g</u>enes <u>a</u>ssociated with <u>m</u>emory after <u>e</u>xercise or <u>s</u>edentary (cGAMES), and <u>c</u>andidate <u>g</u>enes <u>a</u>ssociated with <u>m</u>emory after <u>s</u>edentary (cGAMS) conditions (Fig. 6B-C and Supplementary Table 8). We then evaluated cGAME and cGAMES groups of genes for the new presence or absence of histone modifications H4K8ac and H3K27me3 after ELE, suggesting addition or removal of these histone marks denoting a permissive (increased gene expression) or repressive (reduced gene expression) chromatin state, respectively (Fig. 6B’-C’ and Supplementary Table 9).

**Figure 6:**
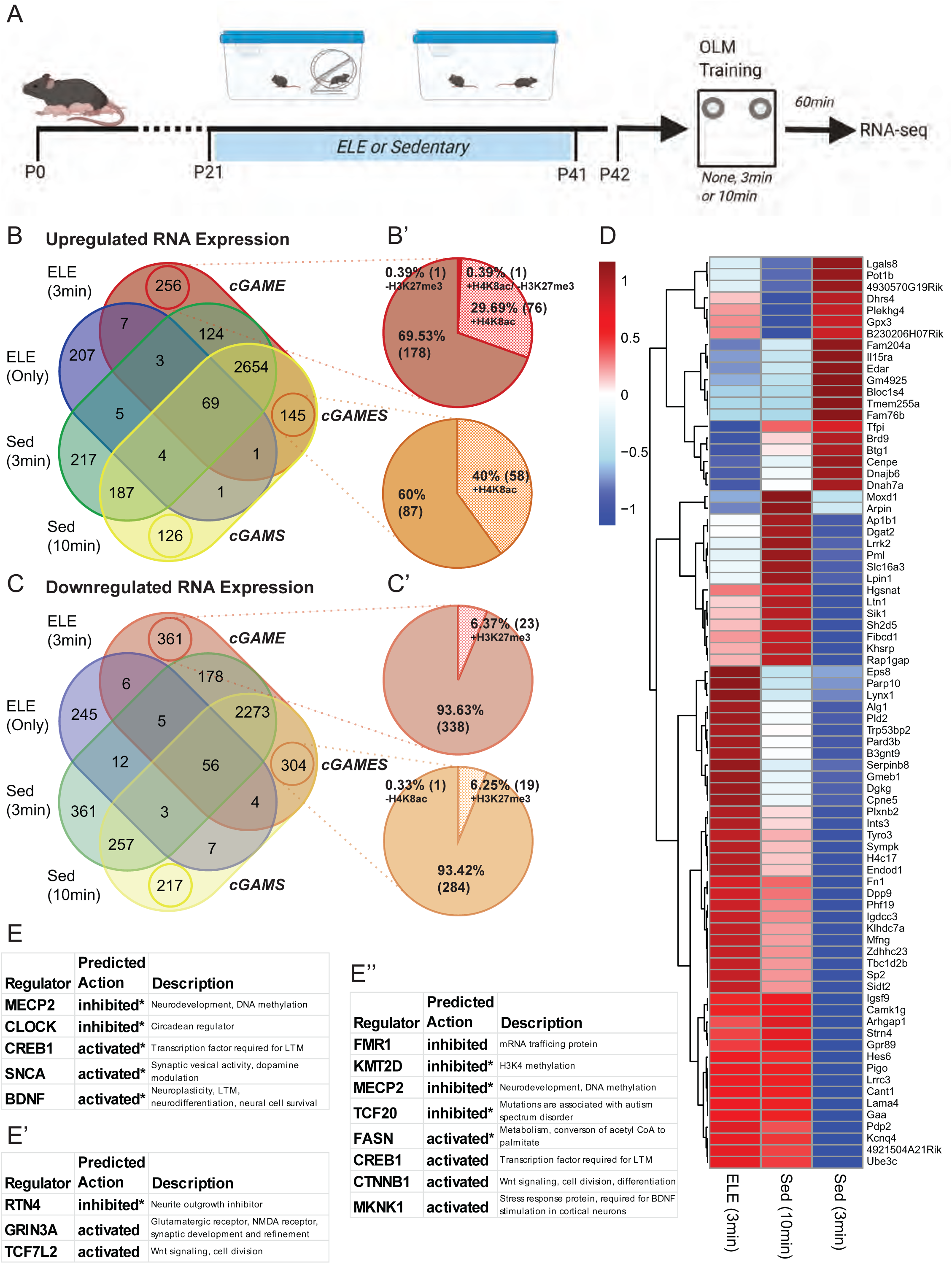
ELE may prime consolidation-induced gene expression. (A) Experimental design diagram. (B) Venn diagram representing upregulated genes relative to mice without ELE or learning in either ELE, ELE and 3 min OLM training, Sed and 3 min OLM training, and Sed and 10 min OLM training. (B’) Pie chart showing the relative percentage of genes that have new H4K8ac presence or H3K27me3 loss as a result of ELE in specific groups identified in the Fig. 6B labeled cGAME and cGAMES. (C) Same as Fig. 6B but for downregulated genes relative to mice without ELE. (C’) Pie chart of downregulated genes in the cGAME or cGAMES groups that have new H3K27me3 or new loss of H4K8ac as a result of ELE. (D) Z-score based heatmap of genes in the cGAMES and cGAME groups that have new H4K8ac or H3K27me3 presence or loss as a result of ELE. Expression values used to calculate the Z-scores were relative to mice that did not experience OLM training or ELE. (E) Upstream regulators identified by IPA of the genes represented by Fig. 6D. (E’) Upstream regulators identified by IPA of the genes in the cGAME (up and downregulated) group that are unique to that when only those are included in the IPA analysis. (E’’) Upstream regulators identified by IPA of the genes in the cGAME and cGAMES (up and downregulated) group when the genes in those groups are put through IPA together. (* indicates that the activated or inhibited indicator is registered from the IPAs based on the z-score for that regulator’s expression and not directly indicated based on software prediction).

In the cGAMES group, there were 145 genes found to be significantly increased in the mice that underwent a threshold learning event (3 min for ELE; 10 min for sedentary mice). Of these 145 genes, 40% of them (58) had new H4K8ac following ELE (Fig. 6 B-B’). We consider this group of genes to potentially be “primed,” or readied, by ELE for rapid transcription to support hippocampal memory consolidation at 3 min OLM training, while also being involved in memory consolidation mechanisms in the threshold learning of sedentary mice (10 min OLM training). The cGAME group of genes are those which significantly increased in the ELE mice that underwent a threshold learning event but were not increased in any of the other groups (including the sedentary, threshold learning event / cGAMS group). These genes may be of unique importance for memory consolidation specifically occurring after ELE given their absence in sedentary learners. We found that the cGAME group had significant upregulation of 256 genes, with approximately 30% (76) of those genes receiving new H4K8ac following ELE alone (Fig. 6 B-B’). We also evaluated the cGAMES and cGAME that were significantly downregulated during memory consolidation. We found 304 cGAMES and 361 cGAME genes (Fig. 6 C). In both of these groups, a small minority of down-regulated genes had altered H3K27me3 after ELE, and this was mostly gain of H3K27me3 (cGAMES: 6.3%; cGAME: 6.4%) (Fig. 6 C’).

We next wanted to determine how ELE “priming” directed gene expression during memory consolidation that occurs after exercise (ELE 3min OLM) relative to sedentary memory consolidation (Sed 10min OLM). We compared these to the group that experienced the sub-theshold learning event (Sed 3min OLM). We compared the relative log_2_ fold changes (LFCs) of the genes in cGAME and cGAMES using z-scores for each group across the genes (Fig. 6D). In the vast majority of these primed genes, the ELE drove gene expression during 3 min of OLM in the same direction as 10 minutes of OLM training in sedentary animals (Fig. 6D). In some cases, the gene expression changes were more exaggerated in the same direction as a sedentary threshold event (Fig. 6D). We interpret this to indicate that ELE enables a 3min consolidation event to be threshold (when 10 minutes is required for a sedentary mouse) by priming these genes for altered gene expression in the same direction as a sedentary threshold event.

We next identified upstream regulators of cGAME and cGAMES genes with ELE-induced histone modifications, as these mediators are likely involved in long-term memory formation of a subthreshold learning event following ELE. The cGAMES and cGAME lists were evaluated using Qiagen’s Ingenuity Pathway Analysis (IPA). First, candidate regulators were identified for the cGAME and cGAMES groups that have histone PTM changes as a result of ELE that we measured (Fig. 6E and Supplementary Table 10). Inhibition of MECP2 may indicate a reduction in DNA methylation. CREB1 is required for LTM^45, 74^. SNCA may play a role in neuroplasticity by modulating synaptic vesicle transport^75^.

Next, we wanted to identify gene networks that might be at play that are not just regulated by the histone PTMs we selected. Many more histone modifications exist and may regulate gene expression during consolidation in addition to those we selected. We investigated the genes in cGAME and cGAMES using IPA and identified a set of likely upstream regulators for these genes taken together (Fig. 6E’’ and Supplementary Table 10). Once again MECP2 inhibition and CREB1 activation appear to be indicated validating their presence in the previous IPA. Notably new to this group are inhibition of FMR1, KMT2D, and TCF20, along with activation of FASN, CTNNB1, and MKNK1. Next, we interrogated what networks might be unique to exercise-enabled consolidation by running the cGAME list through IPA analysis (Fig. 6E’ and Supplementary Table 10). Unique to this group, RTN4, an inhibitor of neurite outgrowth^76^, was predicted to trend towards inhibition. GRIN3A, a glutamatergic NMDA receptor important to synaptic development and refinement^77^, was predicted activated in this group. These findings suggest that ELE may enable consolidation by increasing neurite and synapse growth and development. TCF7L2 is also predicted to be activated. Given that this transcription factor is involved in Wnt signaling, and neurogenesis^78^, and activated CTNNB1 is also predicted to be associated with threshold consolidation generally, altered Wnt signaling may be critical for ELE enabled consolidation.

## Discussion

Identifying gene regulatory mechanisms activated by exercise during sensitive developmental periods can aid in understanding how early-life exercise may promote a “molecular memory” of exercise^23, 59^, via the epigenome, that informs long-term cell function and behavioral outcomes. Here we present results demonstrating neuron-specific, simultaneous characterization of translating mRNA and associated histone modifications to reveal molecular signatures of the early-life exercise experience. To our knowledge, this is the first report describing use of the NuTRAP construct in a predominantly neuronal population. Additionally, our unique experimental approach for simultaneous INTACT and TRAP (“SIT”) is a technical advance for NuTRAP applications: by modifying the nuclear chromatin isolation (INTACT) procedure to include cycloheximide, one can obtain both polyribosome-bound mRNA (using TRAP) and nuclear DNA from a single lysate to directly correlate gene expression programs with epigenetic regulatory mechanisms in the same set of cells. The field of neuroepigenetics has been challenged by very limited approaches for coupling cell-type specific transcriptional profiles with chromatin modifications without compromising cell integrity (through flow sorting) or genomic coverage (through single-cell sequencing). Therefore, the NuTRAP model in combination with the “SIT” protocol offers an exciting opportunity to overcome these challenges and perform paired transcriptomic and epigenomic analyses in any brain cell type of interest for which a Cre line exists^28, 39^.

This study focuses on the impact of exercise on hippocampal gene expression and associated epigenetic modifications specifically during juvenile-adolescent periods (the fourth through sixth postnatal weeks of life in mice). We discover gene expression programs, and alterations in the histone modifications H4K8ac and H3K27me3, that may be unique to this early-life period. There have been recently published reports investigating the effects of voluntary exercise on adolescent and adult hippocampal gene expression and epigenetic modifications through RNA and DNA sequencing methods from heterogenous cell populations^19, 20^. Our data revealed several notable genes with either chromatin-modifying or neurogenesis-regulating properties were either upregulated with new H4K8ac (*Prox1*, *Sntb2*, and *Kptn*), or downregulated with new H3K27me3 (*Col3a1*, *Efnb2*, *Epop*, and *Myoc*). One could consider the mechanistic implications for these genes: for example, ELE could “prime” the expression of *Prox1* by the addition of H4K8ac, and in turn, subsequent *Prox1* expression can promote neurogenic fate and cell location assignment^79^. ELE was also found to alter the expression of *Kptn,* which encodes a protein involved in actin dynamics and neuromorphogenesis^65^. Increased *Kptn* expression could promote cytoskeletal changes occurring with altered synaptic plasticity after exercise. Overall, the differentially expressed genes and their associated chromatin states occurring after ELE provide molecular candidates involved in the effects of ELE on hippocampal function.

Traditional epigenetic and gene expression sequencing approaches to bulk tissue samples are confounded by the cellular heterogeneity. Further, current methods for paired analysis of the epigenome and transcriptome are restricted by the need for multiple transgenic animal models or the use of highly expensive single-cell sequencing technologies. It is critical to obtain epigenomic information in a cell-type specific manner because there are unique epigenetic modifications distinguishing cell subtypes of the central nervous system^10, 80^. Roh et al. first developed the NuTRAP mouse to characterize adipocyte-specific and hepatocyte-specific epigenetic and transcriptional profiles^29^. The NuTRAP mouse has since been used to perform paired transcriptome and DNA modification analysis in astrocytes and microglia, demonstrating its utility in CNS cell-types^39^. The Emx1-NuTRAP mouse described here provides a rapid, cost-effective means for characterizing neuron-specific changes to epigenetic state and gene expression using whole-tissue homogenates containing multiple cell types. In our technical validation experiments, the “SIT” technique produced high quality RNA and DNA of sufficient concentration for downstream sequencing studies from a single mouse hippocampus. This method effectively increases the total sample size used for each analysis as the tissue sample does not need to be separated for DNA and RNA isolation. CUT&RUN allows for low-input cell-specific chromatin analysis^36, 60^ further reducing the required amount of starting material. Indeed, the DNA obtained from SIT could be used for other applications for evaluating the chromatin landscape, including ATAC-seq. Presently, the field of neuroepigenetics is shifting to single-cell resolution for sequencing studies. Our study was performed in bulk tissue enriched for neurons. As the goal of this work was to characterize the transcriptional and epigenomic effects of early-life exercise, cell subtype identity (as is often a priority in single-cell approaches) was not necessary for our initial hypotheses but would offer a second layer of complexity to address this study’s questions. Single-cell approaches can bias against lowly-expressed genes, which may present the risk of not capturing small expression changes in rate-limiting genes that may have large changes in function as a result of ELE.

In this study, we also assessed for hemispheric differences in DEGs and histone modifications using our SIT protocol. A common approach to paired transcriptomic and epigenomic analyses is to use separate brain tissue samples, taking from one hemisphere for gene expression studies and the other side for epigenetic analyses^81^. A previous study utilizing the NuTRAP allele in non-neuronal brain cells used separate brain hemispheres to perform INTACT and TRAP^39^. However, this assumes that cells of the same type from each half of the tissue will show the same epigenetic and transcriptional profile. This is not necessarily the case for all cell types, especially in the CNS, as structures of the brain can show high degrees of hemispheric specialization^82–86^. To address the degree of transcriptional or epigenetic difference between hippocampi from opposite hemispheres, we performed separate INTACT and TRAP isolations using hippocampal tissue from one hemisphere for DNA isolation and the contralateral hippocampus for mRNA isolation. We found that although DEGs and histone PTMs differed between left and right hippocampi, functional analysis using Panther GO revealed similar biological processes between the two hemispheres. We interpret these data to suggest that hemispheric origin of hippocampal tissue is not a critical contributor to conclusions drawn from these types of sequencing studies.

Transcription is required for both hippocampal memory and LTP. LTP is defined as a synaptic plasticity response considered the cellular correlate of learning and memory^87^. Given our previous finding that ELE can facilitate long-term memory and increase LTP ^17^, we hypothesize that this may be due to altering the epigenomic landscape, to ultimately promote a transcriptional state supporting hippocampal memory consolidation. This hypothesis is supported by the finding that in adult rodents, long-term memory is formed in response to a subthreshold learning event after exercise or HDAC3 inhibition and involves increased acetylation of neuroplasticity gene promoters^22^. This study identifies, for the first time, novel gene expression networks and their epigenetic regulation that may be involved in the hippocampal memory benefits of ELE. We also identified a group of genes that are altered in their gene expression only in sedentary animals undergoing a threshold learning event (“cGAMS” group). The separation of this group from the cGAME and cGAMES groups indicates that ELE may engage a separate, perhaps complementary network of genes to promote memory consolidation. A next step of this work will be to test the necessity and sufficiency of identified genes and gene networks, their histone modifications, and/or their upstream regulators mediating the effects of ELE on hippocampal memory performance. Another unanswered question is how long the memory-promoting effects of ELE last after exercise cessation, and whether there is stability of ELE-induced epigenetic modifications to allow for the persistence of ELE effects.

In summary, our work reveals ELE-induced transcriptomic and epigenomic signatures in hippocampal neurons using the Emx1-NuTRAP mouse for such comparisons. A limitation for the use of the NuTRAP transgenic mouse is the requirement of a Cre line to exist for the experimenter’s cell type of interest. In our Emx1-NuTRAP model, the Cre-recombinase in Emx1 expressing cells is constitutively expressed, and this may present off-target effects when present during development. Despite this, our study uses a feasible, previously undescribed approach in the field of neuroepigenetics to pair transcriptomic and epigenomic sequencing results from a population of hippocampal neurons. Our approach revealed new molecular epigenetic candidates for further mechanistic studies to examine the effects of early-life exercise on hippocampal function. Furthermore, understanding how exercise uniquely impacts neuronal function during juvenile-adolescent developmental periods could provide insights into an early-life epigenetic memory of the ELE experience, revealing possible targets for improving memory function or mitigating memory deficits in the setting of early-life cognitive and neurodevelopmental disorders^88^.

## Acknowledgements

This work was supported by the National Institute of Neurological Disorders and Stroke of the National Institutes of Health, under award number K12NS098482, the Conte Center @ UCI Seed Grant, UC Irvine Institute for Clinical and Translational Sciences Pilot Award, and the Robert Wood Johnson Foundation Amos Medical Faculty Development Program (awarded to A.S.I.). We thank the UC Irvine School of Medicine, Departments of Pediatrics for support. We would also like to thank Dr. Marcelo Wood for scientific discussions and critical input on the project, and Drs. Melanie Oakes and Jenny Wu of the UC Irvine Genomics High Throughput Sequencing Facility for next generation sequencing experiments support.

## Author Contributions

A.M.R., T.D.F., N.E.N. and A.S.I. designed the studies and interpreted the results. A.M.R., T.D.F., and A.S.I. performed INTACT and TRAP studies. A.M.R. built libraries for RNA- and TRAP-sequencing. N.E.N. performed CUT&RUN-seq experiments and built libraries for sequencing. N.E.N. and A.S.I. performed flow cytometry experiments. A.M.R. and A.S.I. performed immunostainings and immunofluorescent imaging. D.A.V. and A.B. built exercise cages and monitored running behavior. A.M.R. performed all bioinformatic analysis of sequencing data. A.M.R. and N.E.N. analyzed sequencing data, performed GO, GSEA and IPA analyses, and all other downstream analyses except where noted. A.S.I. performed behavioral studies. T.D.F. performed Spearman’s correlations and gene length by fold change analysis. T.D.F. performed qPCR. N.E.N. designed the figures. A.M.R., T.D.F., and A.S.I. wrote and edited the manuscript, with N.E.N. providing edits to the manuscript. All authors discussed the results and edited and approved the manuscript.

## Methods

### Resource Availability

#### Lead Contact

Further information and requests for resources and reagents should be directed to and will be fulfilled by the lead contact, Dr. Autumn Ivy

#### Materials Availability

- No new reagents or materials were generated for this study.
- The mouse lines used for this study are available from Jackson Labs (Emx1-IRES-*Cre* knock-in mice, Jackson Laboratory Stock No: 005628, and NuTRAP mice, Jackson Laboratory Stock No: 029899).

#### Animals

Emx1-IRES-*Cre* knock-in mice (Jackson Laboratory Stock No: 005628), NuTRAP mice (Jackson Laboratory Stock No: 029899), and C57Bl6/J (Jackson Laboratory Stock No: 000664) wild type mice were obtained from Jackson Laboratories. Female Emx1-IRES-*Cre* and male NuTRAP mice were crossed to generate the Emx1-NuTRAP mice used for this study. Mice were given free access to food and water, and lights were maintained on a standard 12-hour light/dark cycle. On postnatal day (P) 21, mice were weaned and pair-housed in either standard bedding cages or cages equipped with a running wheel. Only male mice were used for the studies performed. Experiments were conducted according to US National Institutes of Health guidelines for animal care and use and were approved by the Institutional Animal Care and Use Committee of the University of California, Irvine.

#### Behavior

Wild type mice were habituated, handled and then underwent OLM training^89^ for either three or ten minutes at P41 as described previously^17^. On postnatal day 36, mice were brought into a testing room with reduced room brightness. Mice were handled for approximately 2 min each, for a total of 5 consecutive days prior to the OLM training session (twice a day for the first 2 days followed by once a day for the next 3 days). Habituation sessions were 5 min, twice per day for 3 days (P39-P41) and occurred within chambers containing four unique spatial cues on each wall of the chamber (horizontal lines, black X, vertical strip, and blank wall). Habituation overlapped with the last 3 days of handling. During the training phase (P42), mice were placed into the same chambers with two identical objects and exposed to either a subthreshold (3 min) or threshold (10 min) training period. Mice were sacrificed for consolidation experiments 1 hour later.

#### Exercise Paradigm

After weaning on P21, male mice from the same litters were randomly divided and pair-housed in either standard cages without a wheel or cages equipped with a stainless-steel running wheel similar to methods already described^17^. Animals housed with running wheels had free access to the wheels from P21–41. Real-time data acquisition (Vital View Software) was used to track distance ran recorded by magnetic sensors that detect wheel revolutions. Each monitored wheel tracked running distance for the entire cage (two mice per cage). Revolutions were quantified for every minute daily for the duration of the exercise period and converted to distance ran per cage (km). Exercised mice that underwent OLM training were individually tracked as described in Valientes et al. 2021^90^.

#### Simultaneous INTACT and TRAP (“SIT”)

On P42 both ELE and sedentary control (n=4/group) Emx1-NuTRAP mice were sacrificed by rapid cervical dislocation and both hemispheres of the hippocampus were dissected out and collected for tissue processing. Hippocampal tissue samples were mechanically homogenized in 1mL of nuclear preparation buffer (NPB) (10mM HEPES (pH7.5); 1.5mM MgCl2; 10mM KCl; 250mM sucrose; 0.1% IGEPAL CA-630; 0.2mM DTT; 100μg/mL cycloheximide; 1x Complete EDTA-free protease inhibitor (Roche)) by razor blade. Samples were then triturated with an 18G needle and syringe 50x on ice.

Homogenized tissue samples were pipetted slowly onto 100μm cell strainer caps on 5mL tubes and centrifuged at 1,000rcf for 10 minutes at 4°C. Cell pellets were resuspended, and samples were pipetted onto 40μm cell strainer caps on 5mL tubes and centrifuged at 1,000rcf for 10 minutes at 4°C. For nuclear isolation, 100μL of Streptavidin-coated Dynabeads were washed twice with 1mL NPB and resuspended with 100μL NPB. Cell pellets were resuspended and added to washed Streptavidin-coated Dynabeads. Samples were briefly pipetted to thoroughly suspend the beads and incubated on ice for 20 minutes. Samples were placed on a magnetic stand to pull down nuclei bound to the beads. The supernatant from each sample was slowly pipetted off the beads and transferred to new 1.5mL tubes for RNA isolation (described later). Nuclei-bound beads were washed twice with 1mL INTACT buffer (10mM HEPES (pH7.5); 1.5mM MgCl2; 10mM KCl; 250mM sucrose; 0.1% IGEPAL CA-630; 1x Complete EDTA-free protease inhibitor (Roche)) and resuspended in 100μL INTACT buffer. Bead-bound nuclei were flash frozen on dry ice and stored at −80°C until ready for CUT&RUN chromatin digestion and DNA purification.

For RNA isolation, 100μL of concentrated TRAP isolation buffer (10mM HEPES (pH 7.5); 117mM MgCl2; 1M KCl; 10% IGEPAL CA-630; 100μg/mL cycloheximide; 11mg/mL sodium heparin; 20mM DTT; 2.2units/μL Rnasin; 1x Complete EDTA-free protease inhibitor (Roche)) was added to each sample (supernatant above). Samples were mixed by gentle pipetting and centrifuged at 16,000rcf for 10 minutes at 4°C. The supernatant from each sample was transferred to new 1.5mL tubes and 50μL of protein-G Dynabeads was added to each sample. (To bind anti-GFP, 100μL of protein-G Dynabeads were washed twice with 500μL NPB and resuspended to a final 100μL volume. 2μL anti-GFP antibody (Sigma-Aldrich, G6539, Lot:128M4867V) was added to each tube and incubated for 1 hour on a rotating nutator at 4°C. Excess anti-GFP was washed off with 200μL NPB and beads were resuspended in a final 100μL volume.) Samples were incubated for 3 hours on a rotating nutator at 4°C. Beads were washed 3x in 1mL high salt wash buffer (HSWB) (50mM Tris (pH 7.5); 12mM MgCl2; 300mM KCl; 1% IGEPAL CA-630; 2mM DTT; 100μg/mL cycloheximide) at 4°C. After the third wash the beads were slowly resuspended in 350µL RLT buffer (from Qiagen RNeasy Mini kit #74004) by slowly pipetting up and down and incubated on a rotating nutator for 30 minutes at room temperature. Samples were placed on a magnetic stand to pull down the beads, and the supernatant from each were transferred to new 1.5mL tubes and stored at −80°C until ready for RNA purification.

#### Separate TRAP RNA Isolation

On P42, mice (n=3 mice/group) were sacrificed and the hippocampus was dissected out as described above. Either the left or right brain hemisphere of the hippocampus was processed for TRAP RNA isolation using methods similar to those previously described^28, 91^ and the other hemisphere was processed for INTACT nuclear isolation (see below). Hippocampal tissue was homogenized with a motorized pestle in 1ml of TRAP homogenization buffer (50mM Tris, (pH 7.5); 12mM MgCl_2_; 100mM KCl; 1% NP-40; 100μg/ml cycloheximide; 1mg/ml sodium heparin; 2mM DTT; 0.2units/μl RNasin; 1x Complete EDTA-free protease inhibitor (Roche)). Samples were centrifuged at 16,000rcf for 10 minutes at 4°C. The supernatant from each sample was collected and incubated with 50μL of protein G Dynabeads (washed twice in 1mL TRAP homogenization buffer without RNasin or protease inhibitor and resuspended in 50μL of the same buffer). Samples were incubated for 30 minutes on a rotating nutator at 4°C. Following the incubation, 100μL of anti-GFP bound protein-G Dynabeads was added to each sample. (To bind anti-GFP, 100μL of protein-G Dynabeads were washed twice with 500μL TRAP homogenization buffer without RNasin or protease inhibitor and resuspended to a final 100μL volume. 2μL anti-GFP antibody (Sigma-Aldrich, G6539, Lot:128M4867V) was added to each tube and incubated for 1 hour on a rotating nutator at 4°C. Excess anti-GFP was washed off with 200μL TRAP homogenization buffer without RNasin or protease inhibitor and beads were resuspended in a final 100μL volume.) Samples were incubated for 3 hours on a rotating nutator at 4°C. Beads were washed 3x in 1mL HSWB at 4°C. After the final wash, the beads were resuspended in 350µL RLT buffer (from Qiagen RNeasy Mini kit #74004) and incubated on a rotating nutator for 30 minutes at room temperature. Following the incubation, samples were transferred to new 1.5mL tubes and stored at −80°C until ready for RNA purification.

#### Separate INTACT Nuclear Isolation

The remaining hippocampal hemisphere from each mouse (not used for TRAP RNA isolation above) was processed for INTACT DNA isolation using methods similar to those described^28, 92^. Tissue samples were quickly chopped with a razor blade, added to 1mL of INTACT buffer, and triturated with an 18G needle and syringe 50x on ice. Samples were then added onto 100μm cell strainer caps on 5mL tubes and centrifuged at 1,000rcf for 10 minutes at 4°C. Cell pellets were resuspended and samples were pipetted onto 40μm cell strainer caps and centrifuged at 1,000rcf for 10 minutes at 4°C. 100μL of streptavidin coated Dynabeads (washed twice with 1mL INTACT buffer and resuspended to a final 100 μL volume) were added to each sample, and samples were incubated on ice for 20 minutes. After incubation, nuclei bound beads were washed twice with 1mL and resuspended in 100μL INTACT buffer. Bead-bound nuclei were flash frozen on dry ice and stored at −80°C until ready for CUT&RUN chromatin digestion and DNA purification.

#### RNA Purification and Sequencing Library Preparation

RNA was purified using the Qiagen RNeasy kit according to the manufacturer’s protocol. After purification, RNA quantity and quality were checked using the Qubit RNA High Sensitivity assay and Agilent Bioanalyzer’s Eukaryote Total RNA Pico assay, respectively, at the University of California, Irvine Genomics High Throughput Facility (UCI GHTF). Using an input of 15ng of total RNA, mRNA was isolated using NEXTFLEX Poly(A) Beads 2.0, and the mRNA sequencing libraries were generated using the NEXTFLEX Rapid Directional RNA-Seq Kit 2.0 according to the manufacturer’s instructions. Final libraries were sent to the UCI GHTF for Qubit dsDNA High Sensitivity assay and the Agilent Bioanalyzer DNA High Sensitivity assay to determine the quantity and quality of the RNA-seq libraries, respectively. Libraries were sequenced at the UCI GHTF using 100bp paired end reads on a single lane of an Illumina NovaSeq6000 to a minimum sequencing depth of 50 million reads.

#### Dorsal Hippocampal RNA Isolation

P42 wild type male mice were sacrificed by cervical dislocation. Left dorsal hippocampus was extracted by bisecting the isolated hippocampus. The tissue was homogenized and RNA was extracted using the Qiagen RNeasy kit according to the manufacturer’s protocol. Sequencing libraries were prepared from the RNA as described under the section “RNA purification” and the UCI GHTF performed sequencing library preparation, except that ERCC ExFold RNA Spike-In Mixes (ThermoFisher cat. 4465739) were added according to the manufacturer’s instructions.

#### Cleavage Under Target and Release Using Nuclease (CUT&RUN)

For chromatin analysis, CUT&RUN was performed using the EpiCypher CUTANA CUT&RUN protocol with slight modifications similar to what has already been described^36^. Nuclei bound to beads were thawed and pulled down on a magnetic stand. The supernatant was removed and beads were resuspended in 200μL wash buffer (20mM HEPES (pH 7.5); 150mM NaCl; 0.5mM spermidine; 1x Complete EDTA-free protease inhibitor (Roche)). Each sample was divided into three aliquots (50μL for anti-H3K27me3 (Cell Signaling Technologies, C36B11, Lot: 16), 100μL for anti-H4K8ac (Epicypher, 13-0047, Lot: 20202001-11), and 50μL for anti-IgG control (Rabbit IgG Fisher Scientific, 026102)). Beads were pulled down on a magnetic stand, resuspended in 100μL wash buffer and incubated for 10 minutes at room temperature. After incubation the beads were pulled down, the supernatant was removed, and the beads were resuspended in 50μL ice cold antibody buffer (20mM HEPES (pH 7.5); 150mM NaCl; 0.5mM spermidine; 0.001mM digitonin; 2mM EDTA; 1x Complete EDTA-free protease inhibitor (Roche)). 0.5μL of the appropriate antibody was added to each sample, and the samples were mixed by gentle pipetting prior to overnight incubation at 4°C on a nutator.

On the second day, the beads were pulled down on a magnetic stand, the supernatant was removed and the beads were washed twice in 250μL ice cold wash buffer with 0.001% digitonin. After the washes, beads were resuspended in 50μL ice cold wash buffer with 0.001% digitonin, and 2.5μL of CUTANA pAG-MNase was added to each sample and mixed by gentle pipetting. Samples were incubated for 10 minutes at room temperature and beads were pulled down on a magnetic stand. The supernatant was removed, and the beads were washed twice in 250μL ice cold wash buffer with 0.001% digitonin and resuspended in 50μL of the same. 1μL of 100mM CaCl_2_ was added to each sample and mixed by gentle pipetting. Samples were incubated for 2 hours on a nutator at 4°C. Following the incubation, 33μL of stop buffer (340mM NaCl; 20mM EDTA; 4mM EGTA; 50μg/mL RNase A; 50μg/mL Glycogen) and 1.65μL of Spike-In DNA (Cell Signaling Technology, #40366) was added to each sample. Samples were incubated in a preheated thermocycler for 10 minutes at 37°C. After incubation, samples were transferred to new 1.5mL tubes and immediately processed for DNA purification.

#### DNA Purification and Library Preparation

DNA was purified using the Monarch PCR and DNA cleanup kit (New England Biolabs, 13-0041) according to the manufacturer’s instructions. After purification, DNA fragments were used to generate sequencing libraries using the NEXTFLEX Rapid DNA-Seq Kit 2.0 according to the manufacturer’s instructions. Indices were diluted to a final concentration of 1:1,000 before being added to samples, and a total of 13 PCR cycles was used for DNA amplification of the libraries. Final libraries were sent to the UCI GHTF for Qubit dsDNA High Sensitivity assay and the Agilent Bioanalyzer DNA High Sensitivity assay to determine the quantity and quality of the CUT&RUN-seq libraries, respectively. Libraries were sequenced at the UCI GHTF using 100bp paired end reads on a single lane of an Illumina NovaSeq6000 to a minimum sequencing depth of 10 million reads.

#### qPCR

cDNA was generated from TRAP-isolated and total input RNA samples (n=2 samples/group) using the Transcriptor First Strand cDNA Synthesis Kit (Roche) following the manufacturer’s instructions. Relative gene expression for *Aqp4* (F: 5’-ATCCAGCTCGATCTTTTGGA −3’, R: 5’-TGAGCTCCACATCAGGACAG −3’), *Tubb3* (F: 5’-GTCTCTAGCCGCGTGAAGTC −3’, R: 5’-GCAGGTCTGAGTCCCCTACA −3’), *Cd11b* (F: 5’-CCCATGACCTTCCAAGAGAA - 3’, R: 5’-ACACTGGTAGAGGGCACCTG −3’), and *Mog* (F: 5’-AAGAGGCAGCAATGGAGTTG −3’, R: 5’-GACCTGCAGGAGGATCGTAG −3’) was determined by qPCR using FastStart Essential DNA Green Master Mix (Roche, 06402712001) following the manufacturer’s protocol. Cycle counts for mRNA quantification were normalized to *Gapdh* (F: 5’-CGTCCCGTAGACAAAATGGT −3’, R: 5’-GAATTTGCCGTGAGTGGAGT −3’). Quantification was performed using the Pfaffl method ^93^.

#### Immunofluoresence

Emx1-NuTRAP mice were anesthetized intraperitoneally with 50mg/kg of sodium pentobarbital and transfused transcardially with 4% paraformaldehyde (PFA) in 1x PBS. After perfusion, whole brains were dissected out and incubated in 4% PFA overnight at 4°C. After fixation, brains were incubated in 30% sucrose for 72 hours and mounted in optimal cutting temperature (OCT) compound. 50μm coronal sections were sliced through the hippocampus using a microtome, and every tenth section was collected in 1mL of cryoprotectant and stored at −20°C until ready for immunofluorescent labeling. For immunofluorescence, sections were brought up to room temperature and washed 2x in 1x PBS on a nutator for 5 minutes each. Sections were then washed 3x in 1x PBS with 0.3% triton-X. After washes, sections were incubated with mCherry polyclonal antibody (ThermoFisher, PA5-34974, Lot: UG2804409F) at a 1:200 dilution in blocking solution (1x PBS with 1% BSA) overnight at 4°C on a nutator. The following day, sections were washed 3x in 1x PBS. Sections were incubated for two hours at room temperature with goat anti-rabbit Alexa Fluor 594 secondary antibody (Invitrogen, A-11012, Lot: 2119134) at a 1:200 dilution in blocking solution. After incubation, the sections were washed twice in 1x PBS. To label nuclei, sections were incubated with DAPI at a 1:10,000 dilution in 1x PBS for 20 minutes. Sections were then washed 3x in 1x PBS and cover slipped with Vectatshield®. Fluorescent images were acquired using a Keyence BZ-X810 All-in-One Fluorescence microscope using the optical sectioning module.

#### Fluorescence Activated Cell Sorting

Fluorescence activated cell sorting (FACS) was performed to characterize neuronal and astrocytic NuTRAP cassette expression. Whole hippocampal tissue from Emx1-NuTRAP mice was isolated and single-cell suspensions were immunostained for cytometric analysis, as previously described^94^. We used antibodies for THY-1 (OX7) AlexaFluor™ 647 (Santa Cruz Biotechnology, sc-53116 AF647, Lot: F2716; concentration: 1:50) and S100β (Abcam, ab41548, Lot:GR3326165-1; concentration: 1:200) with an AlexaFluor™ 405 goat anti-rabbit IgG (H+L) (Invitrogen, A31556, Lot: 2273716; concentration:1:800) secondary antibody. FACS analysis was performed using a BD FACSAria™ Fusion Flow Cytometer (BD Biosciences) at the University of California, Irvine Stem Cell Core. Samples and single-stain controls were analyzed using the FlowJo v10.8.1 software (BD Biosciences). Samples and controls were positively gated for live cells (SSC-A and FSC-A) and single cells (FSC-H and FSC-A), and negatively gated for autofluorescence (Comp-Alexa Fluor 647-A (Thy1) and Comp-BV421-A (S100β)). Fraction of GFP+ neuronal cells was determined using quartile analysis of Thy-1 and Comp-GFP-A (GFP). Fraction of GFP+ astrocytic cells was determined using quartile analysis of S100β and Comp-GFP-A (GFP).

#### Bioinformatics Analysis

##### Reference assembly

For all sequencing analysis that required it, the primary reference assembly used was Genome Reference Consortium Mouse Build 38 patch release 6 (GRCm38.p6).

##### RNA-seq analysis

FastQ files were quality checked for sequencing errors using FastQC (version 0.11.9)^95^. No files were found to have sufficient quality errors to discount their use. Files were aligned using STAR Aligner (version 2.7.3a)^96^. Duplicate reads were removed using Picard Tools (version 1.87)^97^. SAM Tools was used to convert BAM files to SAM files for use in downstream analysis. FastQC, alignment and duplicate removal were all preformed on the High-Powered Compute Cluster (HPC3) operated by The Research Cyberinfrastructure Center (RCIC) at the University of California, Irvine. R (version 4.1.0)^98^ was used for differentially expressed genes (DEG) analysis. Genomic Alignments (version 1.28.0)^99^ (summarizeOverlaps mode=“IntersectionNotEmpty”, singleEnd=FALSE, ignore.strand=FALSE, fragments=TRUE), Genomic Features (version 1.44.0) (exons by gene), and R SAM Tools (version 2.8.0)^100^ (yieldSize=100000) were used to extract a count matrix and generate a summarized experiment object. DEseq2 (version 1.32.0)^101^ was used to perform a DEG analysis. Ensemble IDs were converted to gene symbols using BiomaRt (version 2.48.1)^102, 103^. PCA plots using the top 2000 genes were generated using ggplot2 (version 3.3.4)^104^. Samples were determined as outliers if the variability between the samples heavily weighted PC1 to those samples. This excluded 2 sedentary samples and 1 ELE sample. Heatmaps were generated using gplots (version 3.1.1)^105^ and RColorBrewer (version1.1-2)^106^. Volcano Plots were generated using ggplot2^104^. Venn diagrams were generated using the Venn diagram tool from Bioinformatics and Evolutionary Genomics^107^. Gene ontology analysis was performed using Panther Classification System (version 16.0)^43^. Genes were also categorized and a leading edge heatmap was also generated using Gene Set Enrichment Analysis (version GSEA 4.1.0)^54^. Upstream regulators were identified from the DEGs upregulated by ELE using Qiagen’s Ingenuity Pathway Analysis (IPA) (Fall 2021 Release)^44^.

##### CUT&RUN-seq analysis

FastQ files were quality checked for sequencing errors using FastQC (version 0.11.9)^95^. No files were found to have sufficient quality errors to discount their use. FastQ files were aligned using Bowtie2 (version 2.4.1)^108^. Significant peaks were called from these aligned files using SEACR, developed in CUT&Tag for efficient epigenomic profiling of small samples and single cells^60, 108^, according to CUT&Tag Data Processing and Analysis Tutorial (updated August 12 2020): https://www.protocols.io/view/cut-amp-tag-data-processing-and-analysis-tutorial-bjk2kky. An FDR higher than 0.1 was considered too high for called peaks. This threshold excluded 4 samples (3 ELE H4K8ac separate isolation samples and 1 ELE H4K8ac SIT sample). Peaks were called using the stringent and 0.01 settings. Significant peaks were annotated using ChIPseeker (version 1.8.6)^109^ using TxDb.Mmusculus.UCSC.mm10.knownGene^110^ as a reference with the following settings: tssRegion = c(−3000, 3000), TxDb = TxDb.Mmusculus.UCSC.mm10.knownGene, level = “transcript”, assignGenomicAnnotation = TRUE, genomicAnnotationPriority = c(“Promoter”, “5UTR”, “3UTR”, “Exon”, “Intron”, “Downstream”, “Intergenic”), annoDb = NULL, addFlankGeneInfo = FALSE, flankDistance = 5000, sameStrand = FALSE, ignoreOverlap = FALSE, ignoreUpstream = FALSE, ignoreDownstream = FALSE, overlap = “TSS”, verbose = TRUE). This approach annotated peaks to the closest gene by distance from the promoter, except if that peak falls within a gene before the distal intergenic region. Ensemble IDs were converted to gene symbols using BiomaRt^102, 103^. Gene lists of peaks present in each condition were used to generate Venn diagrams, as with RNA-seq analysis^107^. Further Venn diagrams were generated comparing DEGs with peak calls. Peaks were visualized using UCSC Genome Browser and Genome Browser in a Box^111–113^.

Normalized counts (for read depth) at binned sections of the DNA as well as RNA-seq gene expression count (normalized for read depth) were compared between simultaneous and separate isolations as well as left and right hemispheres using Spearman’s correlations in R with the package ggpubr (version 0.4.0)^114^. Bar plots were generated using Graph Pad 9 PRISM.

#### Data Availability

RNA-, TRAP- and CUT&RUN-seq data will be deposited and publicly available as of the date of publication. Accession numbers will be listed here pending publication. Any additional information required to reanalyze the data reported in this paper is available from the lead contact upon request.

