## Supplemental Figures for "Early life exercise primes the neural epigenome to facilitate gene expression and hippocampal memory consolidation"

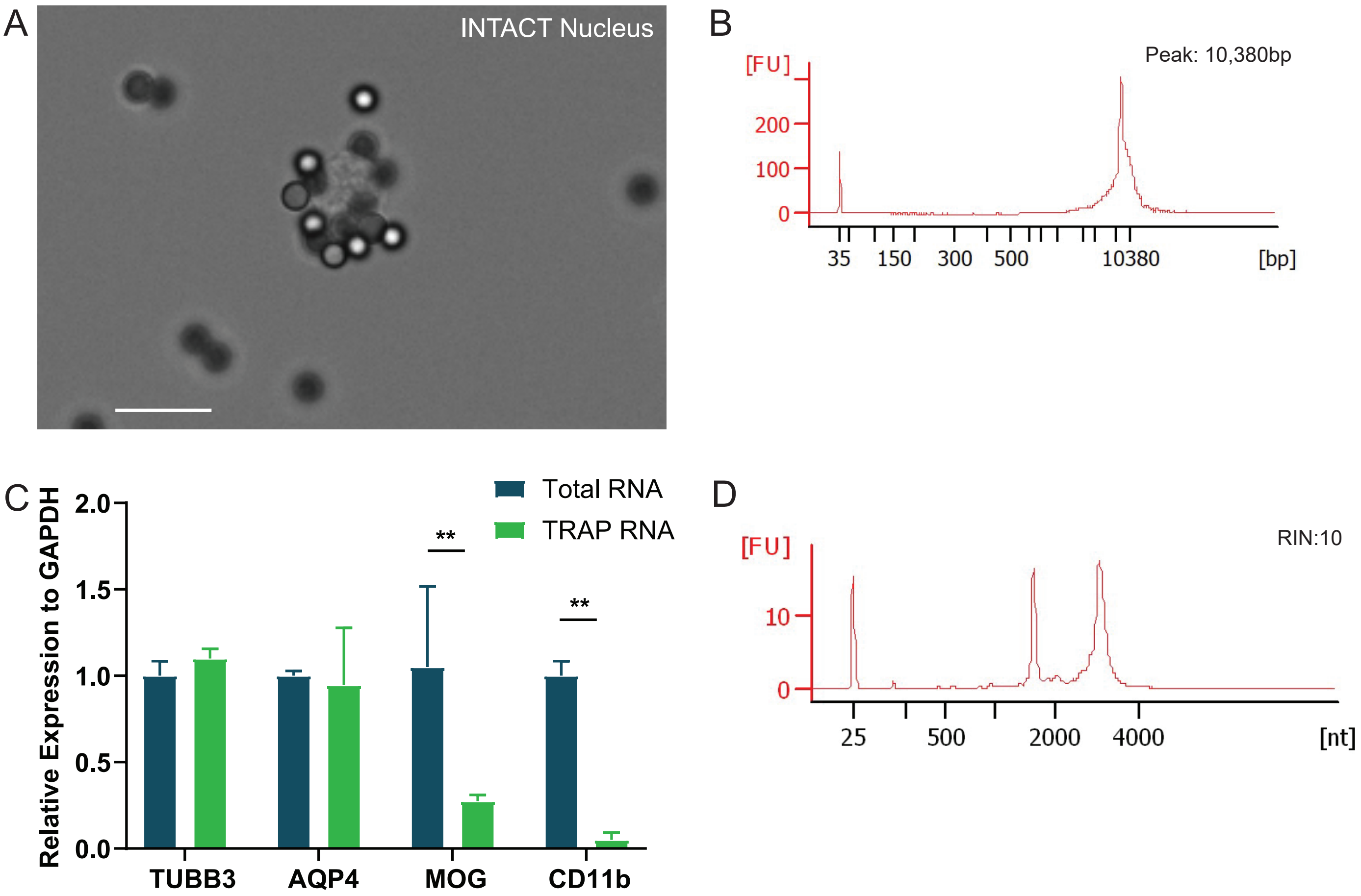
A
INTACT Nucleus
B
Peak: 10,380bp
[FU]
200
100
0
35
150
300
500
10380
[bp]
C
Total RNA
TRAP RNA
Relative Expression to GAPDH
2.0
1.5
1.0
0.5
0.0
**
**
TUBB3
AQP4
MOG
CD11b
D
RIN:10
[FU]
10
0
25
500
2000
4000
[nt]

**Supplementary Figure 1:** (A) Brightfield microscopy of a neural nucleus bound to streptavidin coated Dynabeads™. The scale bar represents 10µm. (B) Representative electropherogram of DNA extracted using the simultaneous isolation protocol. (C) qPCR of TRAP isolated RNA compared to total RNA from the simultaneous isolation protocol **$p<0.01$. (D) Representative electropherogram of RNA extracted using the simultaneous isolation protocol.

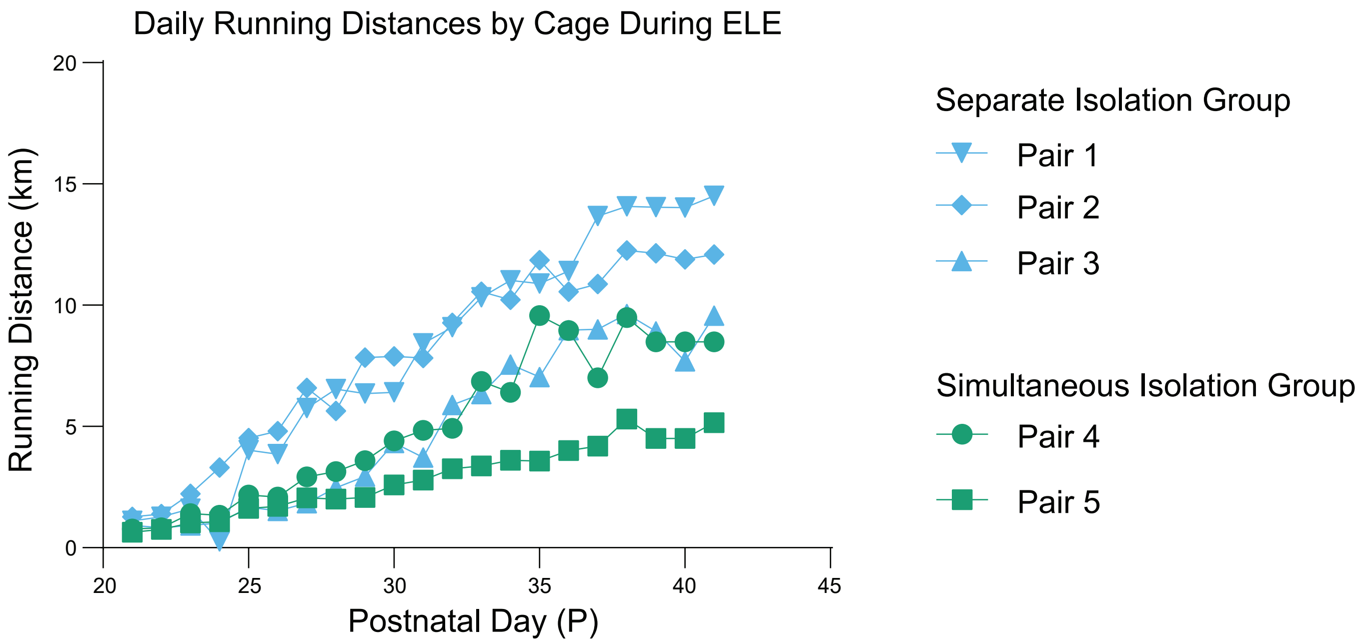

Daily Running Distances by Cage During ELE
Running Distance (km)
Postnatal Day (P)
0
5
10
15
20
20
25
30
35
40
45
Separate Isolation Group
Pair 1
Pair 2
Pair 3
Simultaneous Isolation Group
Pair 4
Pair 5

**Supplementary Figure 2:** Daily running distances of each cage during the ELE paradigm. Values of days missing or low due to software error were imputed by averaging the previous 3 days available. 2 Way Analysis of Variance (ANOVA) reveals that there is significant difference between postnatal days ($p<0.0001$) and cages ($p<0.0001$) but no significant effect of isolation ($p=0.1508$).

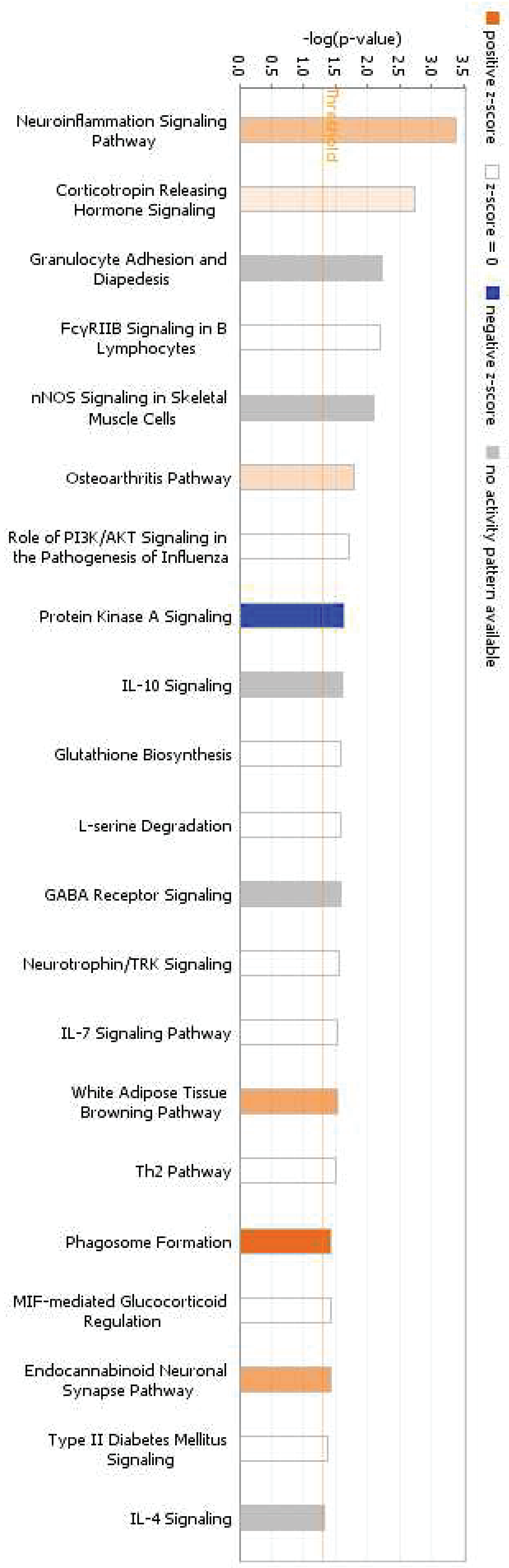


© 2000-2021 QIAGEN. All rights reserved.

**Supplementary Figure 3:** Canonical pathways identified by Qiagen's IPA of genes upregulated by ELE. Supplementary table 3C and 3D have the genes identified by the top two pathways listed in this bar chart.
